# *Mycobacterium tuberculosis* growth arrest on propionate at acidic pH is suppressed by mutations in *phoPR* and pyrazinamide treatment

**DOI:** 10.1101/2025.09.24.678100

**Authors:** Heather M. Murdoch, Shelby J. Dechow, Bassel J. Abdalla, Robert B. Abramovitch

**Author notes:** Co-first authors contributed equally to this study (Heather M. Murdoch, Shelby J. Dechow). **Disclosures**. R.B.A. is an owner of Tarn Biosciences, Inc., a company that is working to develop new antimycobacterial drugs. **Corresponding Author**: Robert B. Abramovitch.

## Abstract

*Mycobacterium tuberculosis* (Mtb) arrests its growth at acidic pH, when grown on specific single carbon sources, including propionate. However, Mtb grows well on propionate at pH 7.0, supporting that propionate can support growth as a sole carbon source. To understand the basis of the propionate-driven growth arrest at acidic pH, we performed a forward genetic selection for mutants that enable growth on propionate at pH 5.7. All the selected mutants had insertions in the two-component regulatory genes *phoR* or *phoP*. We hypothesized that growth arrest at acidic pH is caused by PhoPR diverting carbon from central carbon metabolism towards lipid anabolism and that when PhoPR is inactivated, growth is promoted through metabolizing propionate by the methyl citrate cycle (MCC) into pyruvate, a permissive carbon source for growth at acidic pH. Using chemical inhibition and mutants of the MCC pathway, we demonstrate that the enhanced growth is dependent on the MCC. Furthermore, stimulating lipid synthesis via the methylmalonyl-CoA pathway by adding vitamin B12 restricts growth in the *ΔphoPR* mutant and, conversely, restricting lipid anabolism by inhibiting the triacylglycerol (TAG) synthase *tgs1* enhances growth of the *ΔphoPR* mutant. Notably, CoA pools increased in the *ΔphoPR* mutant grown on propionate, directly supporting our model. Given the role of CoA metabolism in pyrazinamide sensitivity, we examined Mtb sensitivity to pyrazinamide on propionate at acidic pH and, surprisingly, observed that pyrazinamide treatment of WT Mtb suppresses growth arrest on propionate at acidic pH. In contrast, the *phoPR* mutant has enhanced sensitivity to pyrazinamide. Together, these findings support that propionate-driven growth arrest at acidic pH is caused by metabolic remodeling that is regulated by PhoPR and is associated with pyrazinamide sensitivity.

**Importance:** When grown on certain single carbon sources, such as propionate, Mtb arrests its growth at acidic pH and establishes a state of non-replicating persistence (NRP). To understand the genetic basis of this growth restriction, a genetic selection was performed to identify mutants unable to arrest growth at acidic pH with propionate as a sole carbon source. The selection exclusively identified mutants in the PhoPR two-component regulatory system, which functions to modulate cell envelope lipids and redox homeostasis through the upregulation of lipid synthesis at acidic pH. Using genetic and chemical inhibition studies, we demonstrate that PhoPR arrests growth at acidic pH by diverting carbon away from the methyl citrate cycle towards lipid anabolism. Surprisingly, treatment of Mtb with pyrazinamide at acidic pH on propionate, also enabled growth. Therefore, this study defines new mechanisms by which Mtb integrates environmental signaling to regulate growth, metabolism, and drug susceptibility. These findings are relevant to pathogenesis, as PhoPR is essential for growth in macrophages and animals, environments with varying pH and carbon source availability, depending on immune pressures. These data suggest that drug susceptibility may be impacted by enhanced growth and metabolic capacity of Mtb in acidic and propionate-rich environments, such as the within the macrophage or the granuloma.

## Introduction

*Mycobacterium tuberculosis* (Mtb) is a slow-growing pathogen that replicates within macrophages or extracellularly. Depending on the conditions, the bacterium may be colonizing environments with specific carbon sources and varying pH. The combinations of pH, carbon source availability, and other environmental cues can impact Mtb growth rate and virulence. During growth *in vitro*, or *in vivo*, Mtb has a slow doubling time of 20 hours to 70 days(1–3). Under low oxygen, starvation, or mildly acidic pH on specific carbon sources, Mtb enters a non-replicating persistent (NRP) state that allows the pathogen to arrest its growth while maintaining its viability(4). Slow or arrested growth in mycobacteria is thought to play a role in establishing tuberculosis-like diseases and can promote phenotypic drug tolerance and persistence(4–7). Therefore, a mechanistic understanding of how Mtb regulates its growth rate, in response to specific environmental cues, is necessary for understanding pathogenesis and characterizing antibiotic susceptibility.

One of the key regulators of pH-driven adaptations in Mtb is the PhoPR two-component regulatory system (TCS). PhoPR induces the expression of virulence-related genes in response to acidic pH, magnesium, chloride and CO_2_(8–11). These virulence-related genes include the ESX-1 secretion system and cell envelope lipid synthesis genes such as *pks2*, *pks3* and *pks4* which generate sulfolipid (SL) and other acylated trehalose lipids (12, 13). Notably, *phoPR* mutants are highly attenuated in mice and guinea pigs (14, 15), making PhoPR a target for vaccine development(16), supporting that PhoPR and adaptation to acidic pH or related cues are critical for Mtb pathogenesis.

Our lab previously discovered that Mtb has restricted growth on specific single carbon sources at acidic pH. Mtb grows well on carbon sources that function at the intersection of glycolysis and the TCA cycle (e.g. pyruvate, acetate, oleic acid and cholesterol)(17). In contrast, on other tested carbon sources, such as glycerol or propionate, Mtb fully arrests its growth at acidic pH and establishes a state of NRP. We refer to this phenotype as acid growth arrest (17, 18). We hypothesized that Mtb has evolved means to restrict growth at acidic pH on specific carbon sources to regulate growth and support pathogenesis. Indeed, we previously selected for suppressor mutants of acid growth arrest in *ppe51* that enabled Mtb growth on glycerol at acidic pH, increased its replication in infected activated macrophages, and resulted in decreased fitness(18, 19). The goal of this study is to define the mechanisms of acid growth arrest on propionate as a sole carbon source.

Significant research has been conducted examining propionyl-CoA metabolism in Mtb, given it is a catabolic product of cholesterol, a carbon source consumed by Mtb in the host(20). Propionyl-CoA is toxic to Mtb and is detoxified by the methyl citrate cycle (MCC) or by incorporation into methyl branched lipids (such as phthiocerol dimycocerosates (PDIM), SL and acylated trehaloses) (21, 22). The MCC is composed of the methyl-isocitrate lyases *icl1/icl2* along with *prpC* and *prpD* (22). *icl1/icl2, prpC* and *prpD* are essential for growth in minimal media supplemented with propionate (22–25). In the presence of vitamin B12, Mtb is able to survive in the absence of the MCC, by promoting metabolism via methylmalonyl-CoA pathway, enabling the sequestration of propionyl-CoA into methyl branched lipids such as PDIM (21, 22).

To define the mechanism of propionate-driven acid growth arrest, we conducted a forward genetic selection for transposon (Tn) mutants that suppress acid growth arrest on propionate at acidic pH. The selection exclusively identified Tn mutants in the PhoPR TCS. Based on these findings, we propose a model where PhoPR arrests growth by diverting propionate towards the synthesis of cell envelope lipids and, when mutated, growth is promoted by metabolism of propionate by the MCC into pyruvate, a permissive carbon source for growth at acidic pH(17). Additionally, we found that pyrazinamide (PZA) treatment also suppresses the acid growth arrest phenotype in WT Mtb and on propionate PZA selectively sensitizes the *phoPR* mutant to killing PZA. Together, these findings support a model linking acidic pH, carbon source, PhoPR signaling and growth state for PZA activity.

## Results

### PhoPR is required for acid growth arrest on propionate

We previously reported that Mtb grows on propionate at pH 7.0, but arrests growth on at pH 5.7, based on changes in optical density (17). To determine if Mtb is viable and non-replicating, we examined growth by optical density and CFUs over the course of 12 days of Mtb incubated on propionate or pyruvate at pH 5.7. Mtb arrested growth on propionate (Figure 1A) and was viable (Figure 1B), whereas pyruvate was permissive for growth. Therefore, like acid growth arrest on glycerol, Mtb is non-replicating and viable during propionate-induced acid growth arrest.

**Figure 1.**
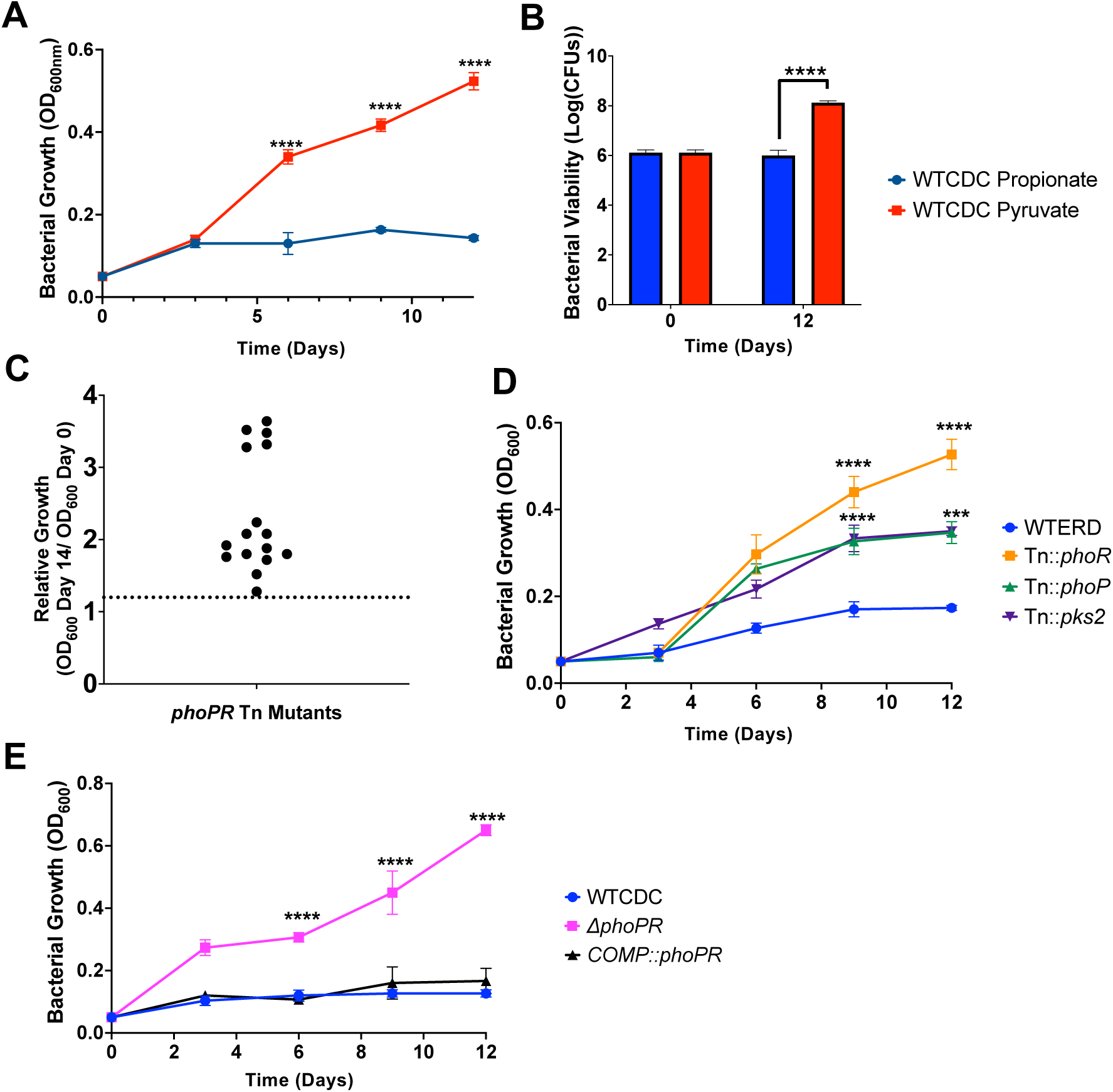
PhoPR arrests growth on propionate at acidic pH. A) Growth curves of WT Mtb CDC strain grown in acidic pH for 12 days, in media supplemented with propionate or pyruvate. B) Bacterial viability evaluated using CFUs plated at days 0 and 12, demonstrating the viability of the growth-arrested bacteria under propionate. C) Relative growth of 16 different transposon mutants with enhanced growth on propionate relative to the WT (dotted line). All sequenced mutants were in *phoR or phoP* (Supplemental Table 1). D) Growth curves of transposon mutants in *phoR*, *phoP* and *pks2*, grown at acidic pH in minimal media supplemented with 2mM propionate. E) Growth curves of *ΔphoPR* mutant and complemented strain compared to WT CDC strain showing growth of the *phoPR* mutant. Multiple comparison unpaired t-test was used for the growth curves analysis while one-way ANOVA was used for the viability analysis, **\*** <0.05, **\*\*** <0.01, **\*\*\*** <0.001, **\*\*\*\*** <0.0001. Experiments were replicated at least twice with similar results.

We hypothesized that growth arrest on propionate is a regulated process and not intrinsic to the carbon source itself, given that the carbon source can be metabolized at neutral pH and Mtb is viable. To test this hypothesis, a forward genetic selection was performed to identify suppressor mutants of acid growth arrest on propionate at acidic pH. An Mtb Erdman transposon (Tn) mutant library containing ∼100,000 mutants was plated on agar plates containing minimal media buffered to pH 5.7 and supplemented with 2mM propionate as a sole carbon source. Mutants with enhanced acidic growth (EAG) phenotype formed colonies on the plates following 8 weeks of incubation. In total, 20 colonies were isolated, and 16 were confirmed as mutants that could grow on propionate (Figure 1C and Table S1). Using whole genome sequencing or inverse PCR, we identified Tn insertion sites in 12 of the 16 confirmed mutants and all sequenced Tn insertions were in *phoR or phoP,* including 6 independent insertions in *phoR and* one insertion in *phoP.* (Supplemental Table 1). Notably, several of the isolated *phoR* mutants also had mutations in the PDIM gene *ppsE*. We confirmed the phenotype of these mutants and observed the *phoR*::Tn and *phoP*::Tn mutants grew on propionate at acidic pH, with higher growth in the *phoR*::Tn mutant (Figure 1D). Using a *ΔphoPR* deletion mutant and complemented strain previously generated in the CDC1551 background (10), we confirmed that at pH 5.7 on propionate the *ΔphoPR* mutant demonstrated robust growth compared to the wildtype and the phenotype was complemented (Figure 1E). There was no significant difference between strains at neutral pH (Figure S1A). Therefore, we conclude that PhoPR is required for acidic growth arrest on propionate at acidic pH.

### The *phoPR* mutant has enhanced growth on multiple carbon sources at acidic pH

Previously, we established that *phoPR* is required to slow the growth of Mtb on the permissive carbon source pyruvate at acidic pH(17). Therefore, we sought to determine if *phoPR* is required for growth arrest on other carbon sources. We observed that the *ΔphoPR* mutant displays enhanced growth on specific carbon sources that are permissive for growth at acidic pH (pyruvate and acetate) and normally non-permissive for growth at acidic pH (propionate and succinate), with the strongest enhanced growth phenotype observed in propionate (Figure 2).

**Figure 2.**
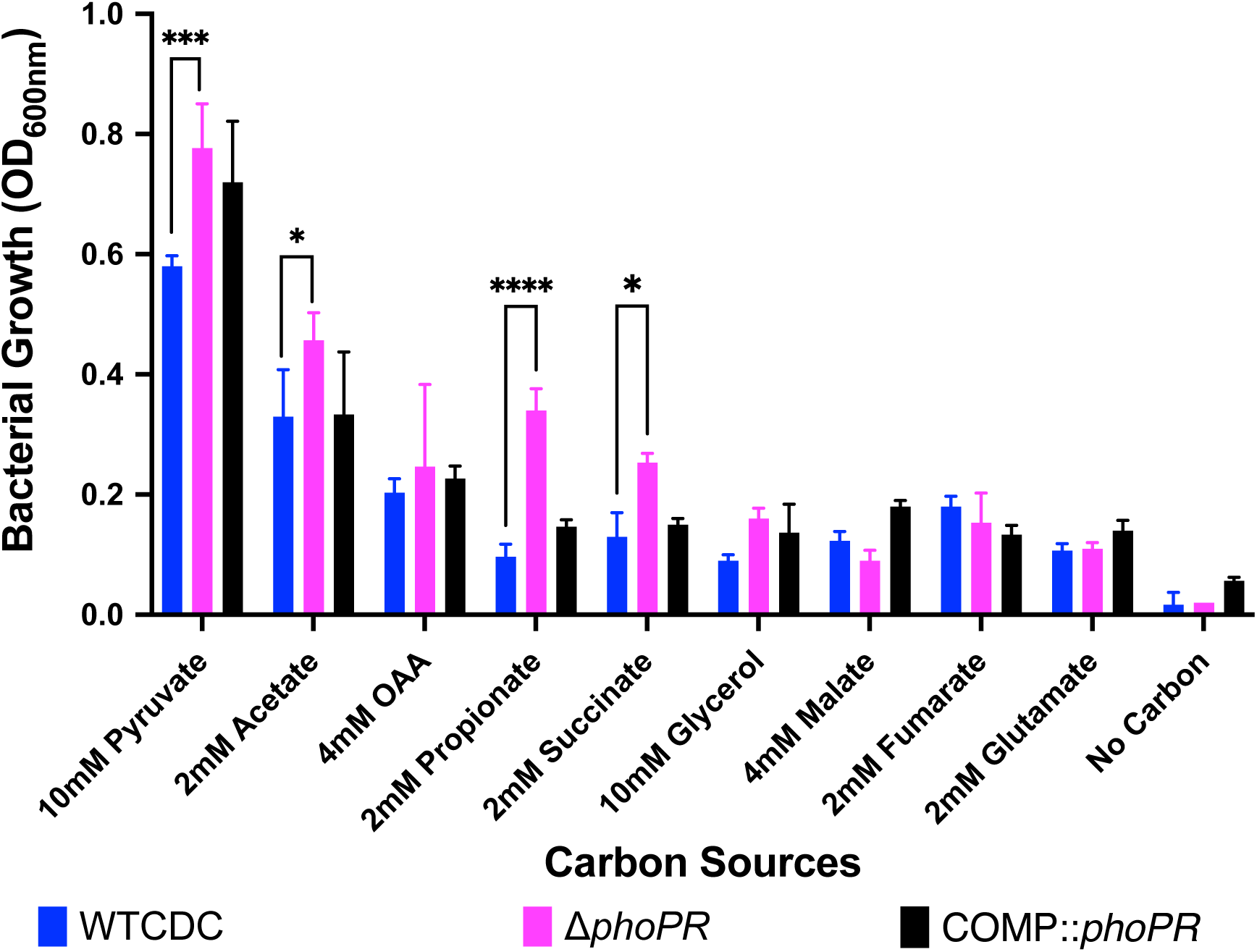
The ΔPhoPR has enhanced growth on specific carbon sources at acidic pH. Mtb growth was evaluated on various carbon sources after 21 days incubation. The *ΔphoPR* mutant exhibits enhanced growth at acidic pH on selected permissive (acetate and pyruvate) and non-permissive carbon sources (propionate and succinate). Two-way ANOVA was used for statistical analysis, **\*** <0.05, **\*\*** <0.01, **\*\*\*** <0.001, **\*\*\*\*** <0.0001. Experiments were replicated at least twice with similar results.

We previously observed that acid growth arrest on glycerol was dependent on both pH and carbon concentration(19). Therefore, we defined the interactions of pH and propionate concentration on Mtb growth by testing growth at varying pH levels (pH 5.0-7.0), and propionate concentrations (0 mM and 20 mM) (Figure 3 A-F). This assay was conducted in 96 well plates where Mtb exhibits less robust growth, as compared to standing flask growth conditions used in prior experiments. At 8 mM and 20 mM propionate, the *phoPR* mutant exhibited enhanced growth at pHs 5.0-5.7, (Figure 3 A-C), whereas at pHs 6.0-7.0, the mutant had similar growth to the wild-type. The *ΔphoPR* mutant exhibited a higher OD at 20 mM propionate, suggesting that growth is proportionate to carbon source availability. Therefore, acid growth arrest is induced at pH 5.7 or below. Notably, the PhoPR regulon is strongly induced pH below 6.0(9), consistent with the induction of PhoPR signaling driving the growth arrest phenotype that we see. *pks2* is strongly induced at acidic pH in a *phoPR-*dependent manner and is required for the synthesis of sulfolipids. At acidic pH, we have previously shown that Mtb promotes SL synthesis and accumulation in a *phoPR*-dependent manner(17). We previously isolated a *pks2::*Tn mutant and hypothesized that if enhanced growth is a consequence of diversion of carbon from the TCA cycle, then the *pks2::Tn* mutant should have enhanced growth on propionate. We examined the growth of the WT,

**Figure 3.**
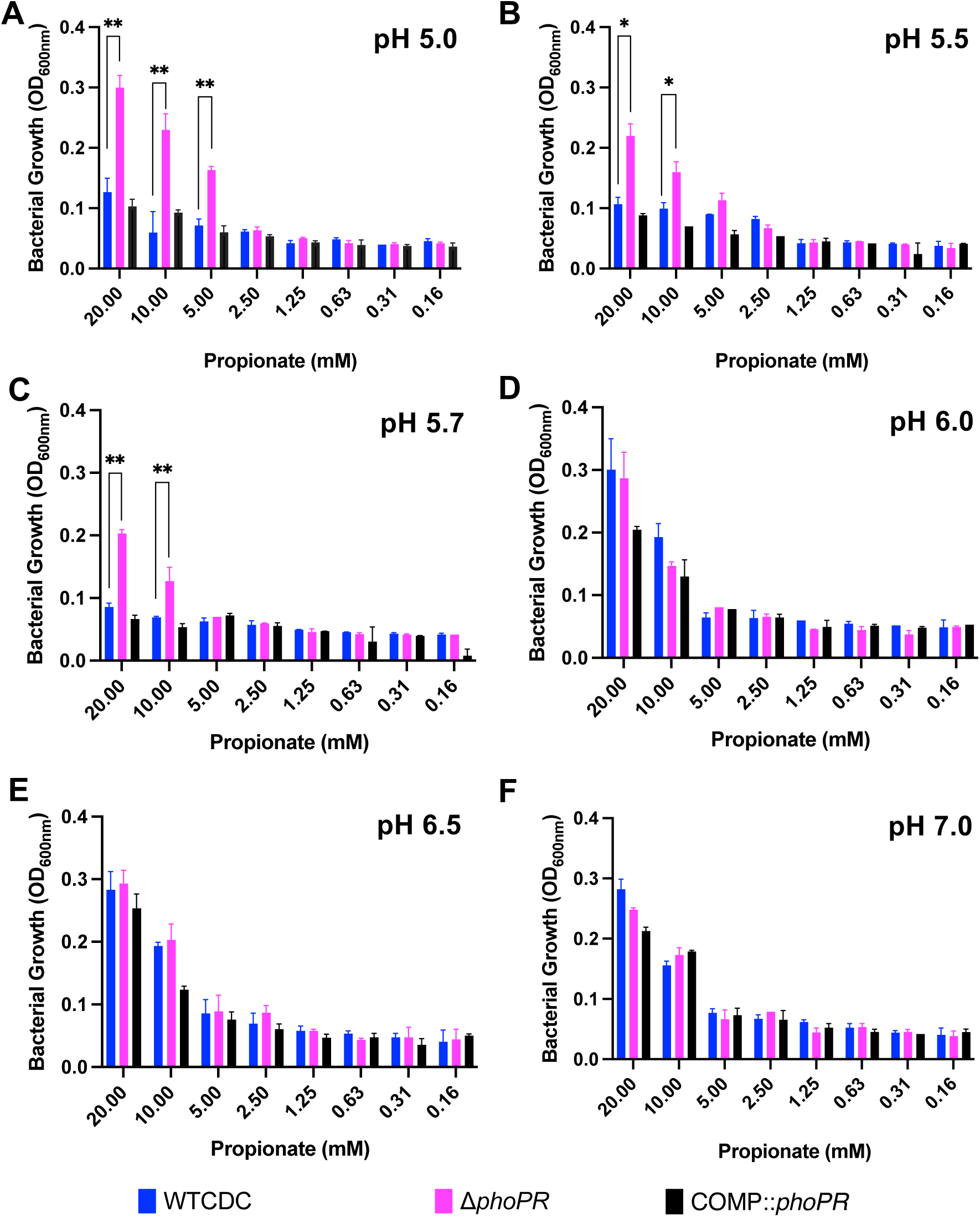
Enhanced growth of the *ΔphoPR* mutant is dependent on level of pH and propionate concentration. Growth of the *ΔphoPR* mutant, the WT and complemented strains was tested in minimal media supplemented with a range of concentrations of propionate (20 mM to 0.16 mM) at different pHs (5.0-7-.0). The *ΔphoPR* mutant has enhanced growth in dependent on propionate concentration at pHs of 5.0, 5.5 and 5.7 (**A-C**), but not 6-7 (**D-F**). A multiple comparison unpaired t-test was used for this analysis, **\*** <0.05, **\*\*** <0.01, **\*\*\*** <0.001, **\*\*\*\*** <0.0001. Experiments were replicated at least twice with similar results.

*phoR*::Tn and *pks2::Tn* mutants on propionate at acidic pH and observed that the *pks2::*Tn mutant displayed an intermediate impact, with higher growth than the WT, but less than that of the *phoR* mutant (Figure 1D). Therefore, the enhanced growth is also dependent on *pks2*, implicating changes in lipid synthesis or the cell envelope composition in growth arrest.

### PhoPR is required for acidic pH-dependent cell death

We were surprised to observe the *phoPR* mutant grow on propionate at pH 5.0 (Figure 3A). This highly acidic pH can lead to cell death. Therefore, we examined WT, *ΔphoPR* mutant and complemented strains for growth and viability over a 12-day time course in 2 mM propionate as a sole carbon source, incubated in standing flasks. As anticipated, we observed a significant reduction of WT OD and CFUs at acidic pH, indicating cell death, and an increase of CFUs for the *ΔphoPR* mutant (Figure 4). Therefore, we conclude that cell death at acidic pH, is not an intrinsic stress associated at acidic pH and instead dependent on PhoPR-dependent activities, possibly related to remodeling of metabolism. Notably, the differences in growth between the pH 5.0 conditions in the dose response study (Figure 3A) and this experiment, are likely driven by the more hypoxic conditions in the 96-well plates as compared to the standing flasks due to differences in head space ratio, which is presumed to result in the more robust growth observed in standing flasks.

**Figure 4.**
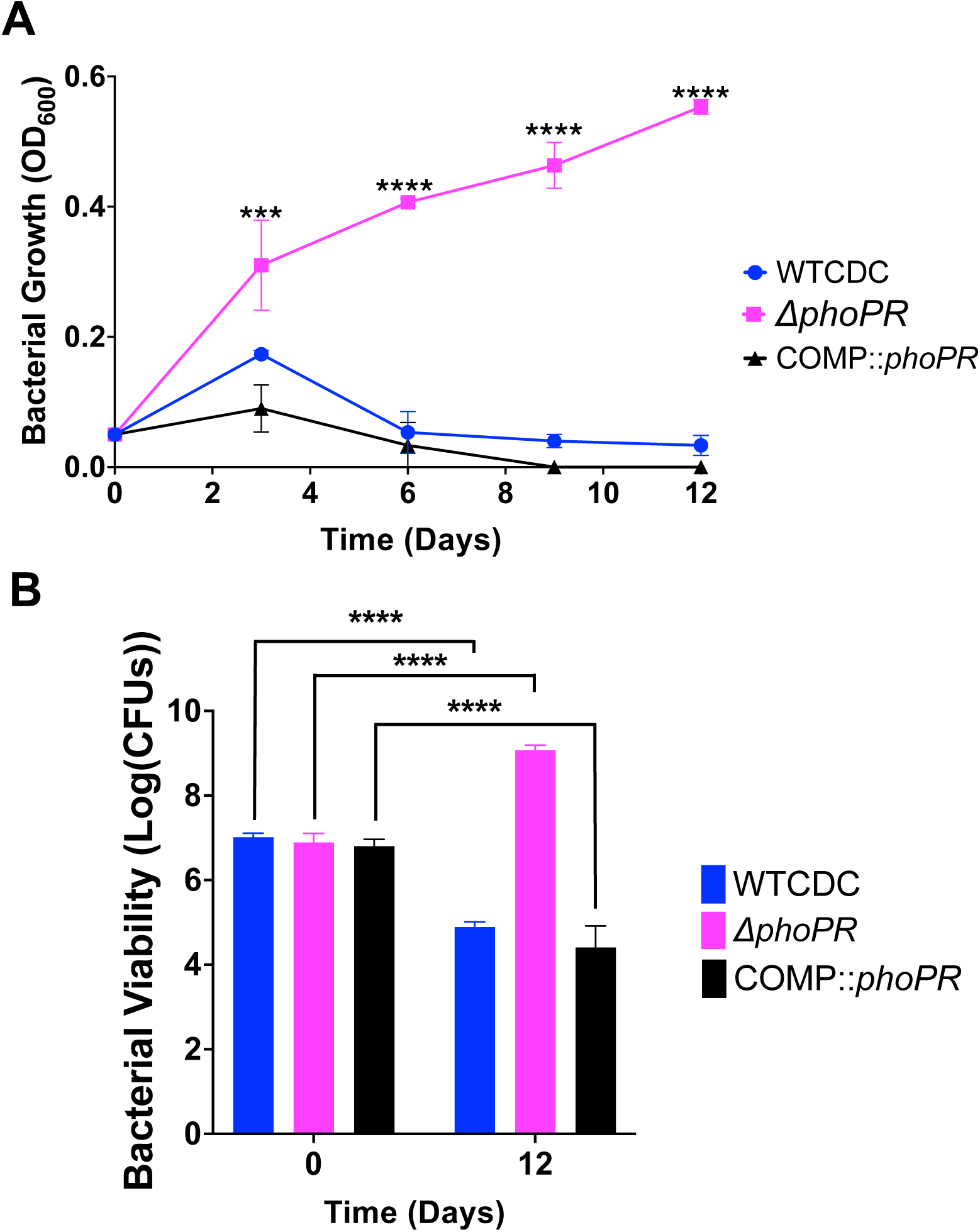
PhoPR is required for cell death at pH 5.0. **A)** The WT, *ΔphoPR* mutant, and complemented strain were incubated at acidic pH for 12 days, in minimal media supplemented with propionate. The *ΔphoPR* mutant has enhanced growth compared to the WT, a phenotype that is complemented. **B)** Viability assays, show that the WT and complemented strains exhibit cell death at pH 5.0, while the *phoPR* mutant grows. A multiple comparison unpaired t-test was used for this analysis, **\*** <0.05, **\*\*** <0.01, **\*\*\*** <0.001, **\*\*\*\*** <0.0001. Experiments were replicated at least twice with similar results.

### PhoPR restricts growth at acidic pH by diverting carbon from the methyl citrate cycle

One of the key functions of PhoPR at acidic pH is envelope composition remodeling by promoting lipid synthesis, including sulfolipids (SL), diacyltrehaloses (DAT) and polyacyltrehaloses (PAT)(13). Deletion of *phoPR* results in a near complete loss of SL at acidic pH, which is compensated for by the induction of triacylglycerol (TAG) (17). Propionyl-CoA is an important substrate for the PhoPR-dependent synthesis of methyl-branched lipids such as SL, DAT and PAT, as well as methyl-branched lipids synthesized independently of PhoPR, such as PDIM and TAG. Given the partially enhanced growth of the *pks2* mutant (Figure 1D), we hypothesized that growth arrest at acidic pH is due to PhoPR diverting carbon away from central metabolism towards lipid metabolism, thus starving Mtb of carbon needed for growth. Indeed, a similar phenomenon has been observed for hypoxia-driven growth arrest and the induction of *tgs1* to produce TAG (26). We further hypothesize that in the PhoPR mutant, the propionate can be metabolized into pyruvate via the methyl citrate cycle (MCC). Pyruvate is permissive for growth at acidic pH, thus the redirection of propionate from lipid anabolism (in the WT) to pyruvate via the MCC (in the *phoPR* mutant) could explain the enhanced growth of the *ΔphoPR* mutant on propionate at acidic pH.

To first test this model, we examined if the MCC was required for growth on propionate at acidic pH, using mutants in the methylisocitrate lyases *icl1/icl2* and complemented strains(24). Lacking a *phoPR* mutant in the *icl1/icl2* mutant backgrounds, we used a chemical inhibitor of *phoPR* signaling, ethoxzolamide (ETZ) (27), to downregulate PhoPR. The strains were cultured in minimal media in propionate with or without ETZ, at acidic pH, and growth and viability were monitored over 12 days(Figure 5A and S2A). In the DMSO control, the WT and *icl1* or *icl2* mutants all had arrested growth at acidic pH, and the *icl1/icl2* exhibited a reduction of CFUs and OD, a phenotype consistent with propionate toxicity in the absence of the MCC (Figure 5A and S2A). In the ETZ-treated cells, we observed enhanced growth of the WT, consistent with PhoPR inhibition driving growth at acidic pH (Figure 5A and S2A). The *icl1* and *icl2* mutants both had reduced growth and viability (Figure 5A and S2A), with a the *icl2* mutant showing a greater reduction in viability which could be driven by the direct regulation of Icl2 by propionate(28). Notably, at pH 7.0, the single *icl1* or *icl2* mutants grew well on propionate in both the DMSO and ETZ treated cells, while the *icl1/icl2* mutant could not grow, due to propionate toxicity (Figure S3). To further confirm the role of the MCC in *phoPR-*dependent enhanced growth, we examined if chemical inhibition of Icl1/2 by itaconic acid (ITA) could inhibit the enhanced growth. Indeed, ITA completely inhibited Mtb *phoPR* mutant growth on propionate at acidic pH (Figure S4). These data are consistent with the MCC being required for enhanced growth of the *ΔphoPR* mutant at acidic pH. To further validate the model that PhoPR-dependent diversion of carbon to lipid synthesis promotes the growth arrest phenotype, we hypothesized that stimulating lipid synthesis in the PhoPR mutant would suppress growth on propionate. To test this hypothesis, we examined whether the induction of the methylmalonyl-CoA pathway, by the supplementation with vitamin B12, reduces the growth of the PhoPR mutant. The WT, *ΔphoPR*, and complement strains were grown in minimal media at acidic pH supplemented with propionate, with or without vitamin B12, and growth and viability were monitored over 12 days. As hypothesized, vitamin B12 suppresses the growth of the *ΔphoPR* mutant at acidic pH (Figure 5B and S2B), supporting that stimulating lipid anabolism suppresses growth at acidic pH on propionate. Addition of vitamin B12 had no impact on Mtb growth at pH 7.0 on propionate , in the WT or *ΔphoPR* mutant (Figure S2D), supporting the phenotypes are acidic pH dependent.

**Figure 5.**
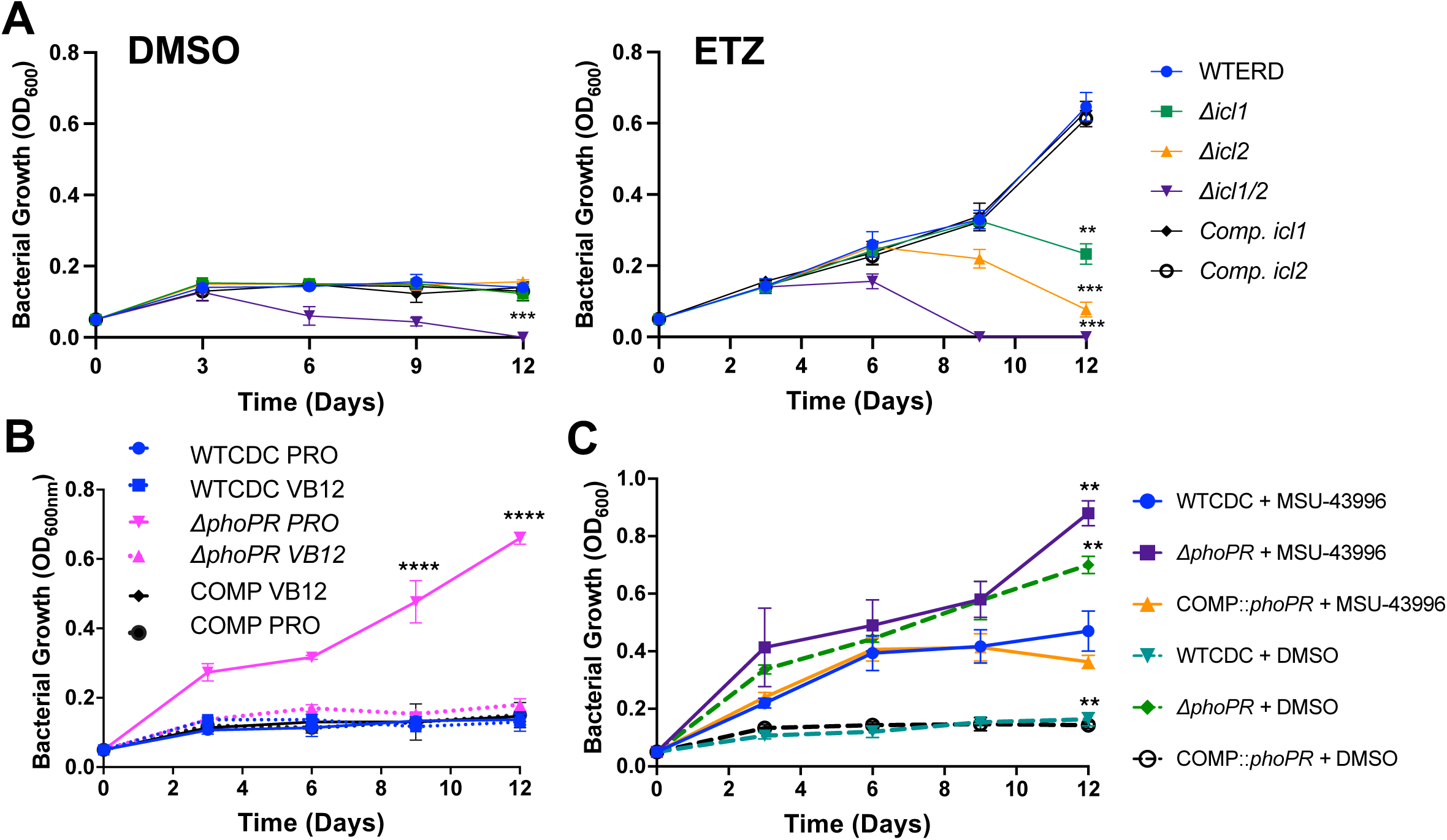
PhoPR restricts growth at acidic pH by diverting carbon from the methyl citrate cycle. **A)** Growth curves of the WT, knockout strain of isocitrate lyases 1 and/or 2, and the complemented strains, treated with DMSO or ethoxzolamide (ETZ), an indirect inhibitor of PhoPR, showing a significant reduction in growth in the *icl2* and double knockout strains. **B)** Growth curves conducted in wildtype, *phoPR* mutant, and complemented strains in minimal media buffered to pH 5.7 supplemented with 2mM propionate and vitamin B12 over a course of 12 days, showing vitamin B12 suppresses enhanced growth in the *ΔphoPR* mutant strain.**C)** Growth of the WT, *ΔphoPR* mutant, and complement strains treated with MSU-43996 or the DMSO control, showing an enhanced growth in the presence of MSU-43996 treatment. An unpaired t-test was used between individual groups and the WT for the growth curves, while a one-way ANOVA was used for the viability experiments, **\*** <0.05, **\*\*** <0.01, **\*\*\*** <0.001, **\*\*\*\*** <0.0001. Experiments were replicated at least twice with similar results.

Mtb increases the production of the TAG when *phoPR* is deleted, possibly as a mechanism to balance redox homeostasis(17). We hypothesized that TAG synthesis may function similarly as a sink for carbon and slow the growth of WT or the *ΔphoPR* mutant. To test this hypothesis, we used a DosRST inhibitor, MSU-43996, that potently inhibits *tgs1* expression and TAG synthesis (29). We examined the growth of WT, and the *ΔphoPR* mutant in the presence of propionate, with or without 40 µM MSU-43996 at acidic pH and neutral pH (Figure 5C and S1B) . When *tgs1* is inhibited, the WT exhibits enhanced growth, and the *ΔphoPR* mutant exhibits further enhanced growth at acidic pH (Figure 5C and S2C); while no major differences are observed at neutral pH (Figure S1B). These data further reinforce the hypothesis that lipid synthesis is slowing or arresting Mtb growth at acidic pH.

### PhoPR restricts CoA at acidic pH

We hypothesize that if PhoPR is diverting carbon away from central metabolism to slow growth, this adaptation may be associated with less available CoA in the WT at acidic pH. In the *ΔphoPR* mutant, metabolism of propionate into pyruvate would fuel the TCA cycle and promote increases in CoA pools. To test this hypothesis, we examined total CoA in the WT, *ΔphoPR* mutant, and complemented strains in 2mM propionate, 10 mM pyruvate or 10 mM glycerol at pH 5.7. During growth arrest in propionate and glycerol, WT CoA pools remained relatively stable over the course of six days (Figures 6A and S5). In contrast, on propionate CoA pools in the *phoPR* mutant increased threefold at day 3 and then lowered again at day 6 (Figure 6A). There is no change between day 0 and 6 in CoA in the growth arrested *phoPR* mutant in the presence of glycerol (Figure S5). In pyruvate, which is growth permissive, CoA pools increased slightly in the WT and were enhanced in the *phoPR* mutant, which also has enhanced growth in pyruvate (Figure 6C). Consistent with our hypothesis, diversion of carbon from the TCA cycle to lipid synthesis by the addition of vitamin B12, decreases the accumulation of CoA in the *phoPR* mutant (Figure 6B), which is also associated with decreased growth (Figure 5B).

**Figure 6:**
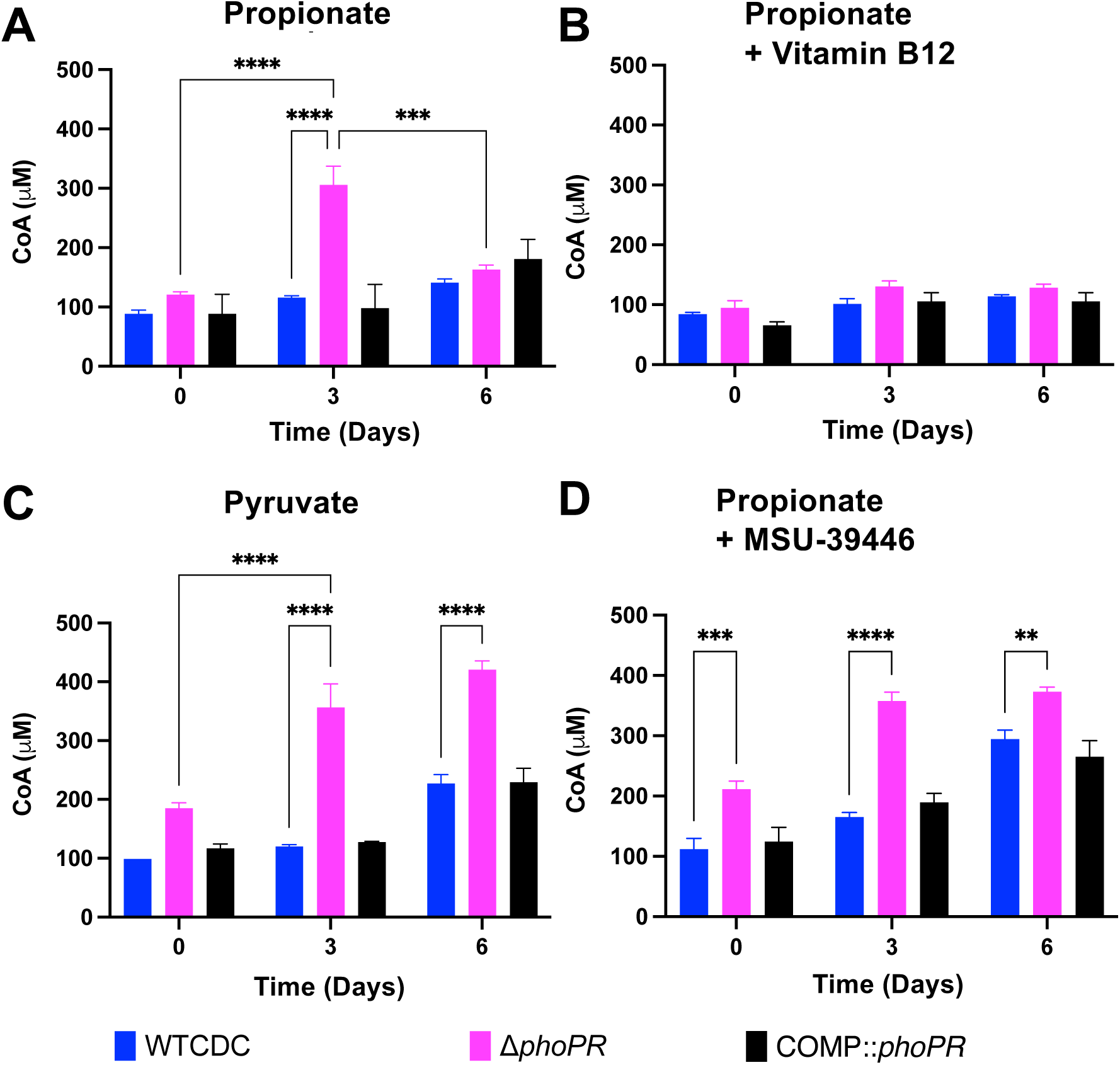
Enhanced acid growth is associated with increased CoA pools. **A)** CoA pools in the *ΔphoPR* mutant, WT CDC, and complemented strain grown in minimal media supplemented with 2 mM propionate at acidic pH, showing enhanced metabolic capacity (i.e., higher CoA pools) in the *ΔphoPR* mutant during the first three days of growth (p = 0.006), which significantly decreases around day 6 (p = 0.009). **B)** CoA pools in the three strains, grown in minimal media supplemented with vitamin B12 at acidic pH, showing no significant differences between the three strains throughout the 6-day growth period. **C)** CoA pools in the three strains, grown in minimal media supplemented with 10 mM of pyruvate at acidic pH, showing enhanced metabolic capacity of the *ΔphoPR* mutant and WT through day 6. **D)** CoA pools in the three strains, grown in minimal media supplemented with 40 µM of MSU-43996 at acidic pH, showing enhanced metabolic capacity of the *ΔphoPR* mutant when *tgs1*, a is inhibited. Two-way ANOVA was used for this analysis, **\*** <0.05, **\*\*** <0.01, **\*\*\*** <0.001, **\*\*\*\*** <0.0001. Experiments were replicated at least twice with similar results.

In our findings, total CoA in the *phoPR* mutant on propionate at acidic pH is negatively associated with stimulation of lipid synthesis pathways and positively associated with growth. The decrease in CoA pools at Day 6 in the *phoPR* mutant suggested that additional lipid synthesis pathways may be engaged over time. The cultures are grown in standing flasks and we anticipate that growth over 6 days by the *phoPR* mutant will consume oxygen and induce the *dosRST* pathway. This would cause the strong induction of the TAG-synthase *tgs1* and provide an induced, second mechanism to restrict growth under the hypoxic conditions caused by the growth of the mutant. To test this hypothesis, we examined the impact of the *dosRST* inhibitor MSU-39446, on CoA pools. In the MSU-39446 treated *phoPR* mutant the CoA pools did not decrease at day 6 (Figure 6D), a phenotype consistent with the enhanced growth caused by MSU-39446 treatment (Figure 5C). Notably, CoA pools increased in the WT, a response consistent with the enhanced growth of the WT (Figure 5C). Together, these data further support the hypothesis that growth arrest on propionate at acidic pH is driven by restriction of central metabolism via synthesis of PhoPR-dependent lipids.

### Pyrazinamide suppresses growth arrest on propionate at acidic pH

Pyrazinamide (PZA) activity is enhanced at acidic pH and its mechanism of action is uncertain, but recent data suggests it may be associated with CoA metabolism and cell envelope lipid homeostasis (17, 30–32). Therefore, we hypothesized that the WT strain and *phoPR* mutant may exhibit differential sensitivity to PZA. To test this hypothesis, we examined the sensitivity of WT, *ΔphoPR* mutant, and complemented strains to PZA across a dose range from 0-100 µM in minimal medium supplemented with 2mM propionate at pH 5.7. Surprisingly, following 21 days of incubation, PZA at concentrations of 2.5 and 10 µM suppressed acid growth arrest in the WT and complemented strains (Figure 7A). However, the *phoPR* mutant was growth arrested under these conditions, showing that it has enhanced sensitivity to PZA, as compared to the DMSO control, where the *phoPR* mutant grew robustly. At a concentration of 1.25 µM, the growth of the WT and *phoPR* mutant are about equal (Figure 7A) ,suggesting a complex interplay between PZA concentration and modulation of growth at acidic pH on propionate.

**Figure 7:**
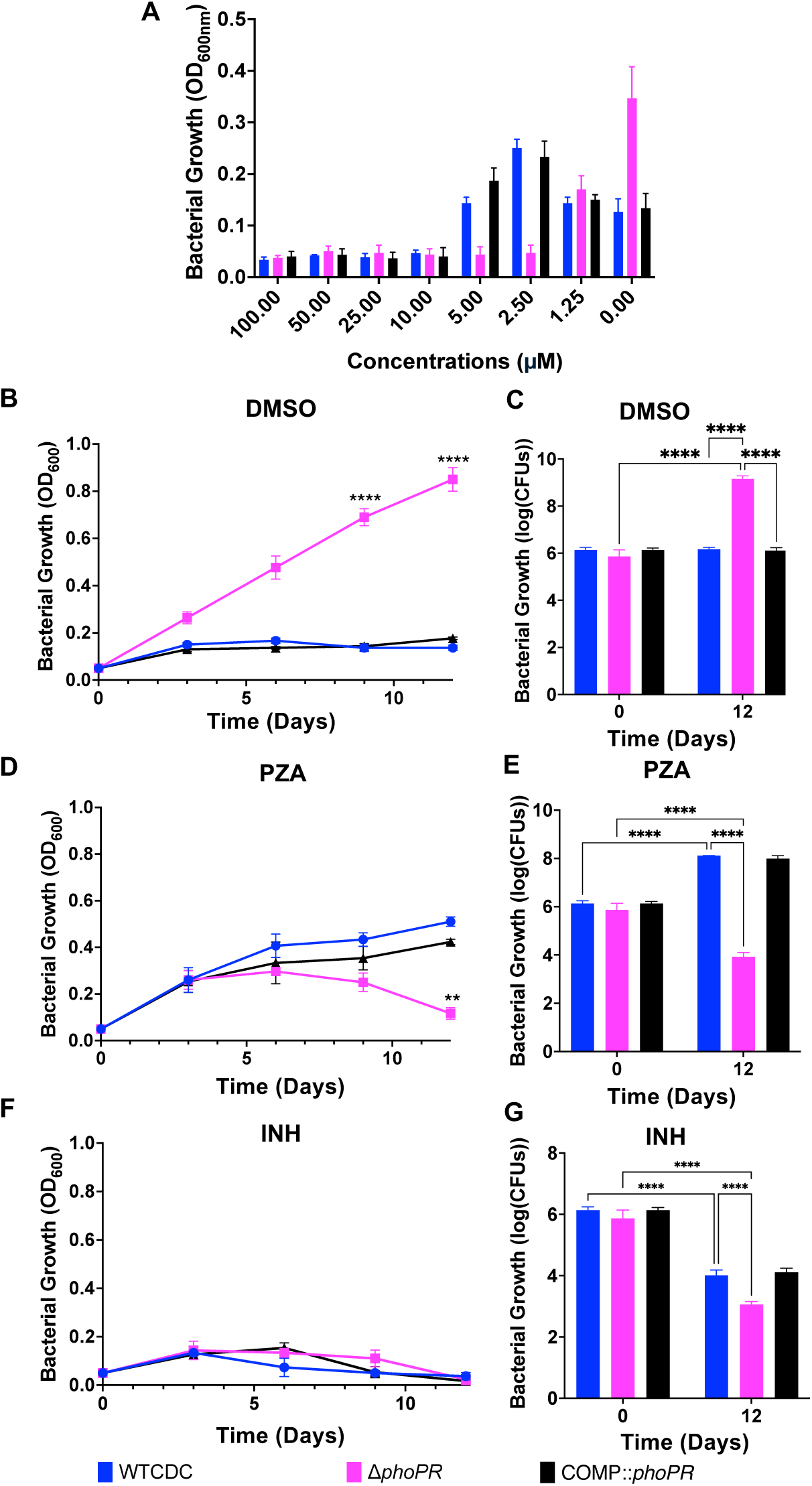
PZA suppresses acid growth arrest on propionate. **A)** Dose-response relationship of pyrazinamide (100-0 µM) in the *ΔphoPR* null mutant, WT, and complement strains, conducted in minimal media supplemented with 2mM propionate at acidic pH, demonstrating enhanced growth of the WT and complemented strains on PZA at concentrations between 5.0-2.5 µM. **B and C)** Assessment of the growth (B) and viability (C) of the three strains in minimal media supplemented with 2mM propionate at acidic pH, treated with the vehicle as a control, showing enhanced growth of the *ΔphoPR* mutant. **D and E)** Assessment of the growth (D) and viability (E) of the three strains in minimal media supplemented with 2mM propionate at acidic pH treated with PZA (3.84 µM), demonstrating enhanced growth of the WT and higher sensitivity of the *ΔphoPR* mutant. **F and G)** Assessment of the growth (F) and viability (G) of the three strains in minimal media supplemented with 2mM propionate at acidic pH treated with isoniazid (INH, 20 µM), demonstrating similar levels of sensitivity in the three strains. Multiple comparison T-test was used for the growth curves analysis, while Two-way ANOVA was used for the viability, **\*** <0.05, **\*\*** <0.01, **\*\*\*** <0.001, **\*\*\*\*** <0.0001. Experiments were replicated at least twice with similar results.

The above assay was conducted in 96-well plates, and we sought to better characterize the suppression of acid growth arrest by PZA in a standing flask-based acid growth arrest assay, which exhibits more robust growth. WT, *ΔphoPR* mutant and complemented strains were incubated at pH 5.7 with propionate as a sole carbon source, treated with 3.8 µM PZA, 20 µM isoniazid (INH), or a DMSO control, and growth was monitored over 12 days. The WT and complemented strains exhibited enhanced growth on PZA as compared to DMSO (Figure 7A and 7B), whereas the *phoPR* mutant was killed by PZA and showed robust growth in the DMSO control (Figure 7D and 7E). We hypothesized that like enhanced growth of the *ΔphoPR* mutant, PZA may also be causing enhanced growth by inhibiting lipid synthesis. To test this hypothesis, we examined if additional of vitamin B12, limited the enhanced growth of PZA treated WT Mtb. As hypothesized, vitamin B12 blocked the enhanced growth in the PZA treated Mtb (Figures S6 and S7). INH inhibited the growth of all of the strains, showing the differential growth impact is PZA-specific (Figure 7F and 7G). We also examined the impact of PZA sensitivity on the strains grown on pyruvate and observed that that WT Mtb did not show enhanced growth on PZA, showing the enhanced growth is propionate specific. (Figure S8). Notably, the *ΔphoPR* mutant was also more sensitive to PZA on pyruvate (Figure S8) and INH on propionate (Fig. 7G), supporting that the enhanced PZA sensitivity is not carbon source specific and the *phoPR* mutant is more sensitive to distinct classes of antibiotics. To determine if the enhanced growth on WT Mtb on PZA is shared between different strains, we also conducted this experiment in Mtb Erdman. We observed that 3.8 µM PZA caused enhanced growth of WT Erdman (Figure S9). No significant differences were observed, between strains or treatments, when the experiment was replicated at neutral pH (Figure S10), an expected result given the acidic pH-dependent activity of PZA. Together, these findings show that PZA can result in enhanced growth or killing of Mtb depending on the pH, concentration of PZA and PhoPR-signaling of the bacterium. This discovery generates further complexity for understanding the mechanism of PZA function, as pH and carbon sources can vary in different environments during infection, and differences in *phoPR* or *prpR* genotype and function exist in clinical strains (33–35).

## Discussion

Mtb remodels its physiology at acidic pH below ∼6.5, demonstrating several pH-dependent adaptations associated with pathogenesis and drug susceptibility(5, 36). Changes in carbon metabolism are a substantial component of acidic pH-dependent adaptations. We and others have observed that Mtb restricts its ability to grow on specific carbon sources at acidic pH, a phenomenon we refer to as acid growth arrest. Previously, we found that Mtb arrests its growth on glycerol and that this growth arrest is overcome by mutations *in ppe51* that promote enhanced glycerol uptake (36). We concluded that growth arrest is overcome by increasing glycerol uptake (36), possibly to overcome reduced glyceraldehyde-3-phosphate dehydrogenase activity at acidic pH(37). *ppe51* mutants did not promote growth on other non-permissive carbon sources, including propionate, leading us to hypothesize that acid growth arrest has different mechanisms for different carbon sources.

The isolation of *phoPR* mutants that can grow on propionate strongly support our hypothesis of carbon source specific metabolic restraints at acidic pH (Figure 8). For example, *phoPR* mutants cannot grow on glycerol and *ppe51* EAG variants cannot grow on propionate. Together, multiple independent observations support that the growth arrest on propionate is due to the diversion of carbon towards the synthesis of PhoPR-dependent lipids (Figure 8). For example, we observe growth arrest at acidic pH when methyl-branched lipid synthesis pathways are activated on propionate, and when PhoPR-dependent pathways (SL, DAT and PAT), DosRST-dependent pathways (TAG), or vitamin B12-dependent pathways (PDIM) are available to the pathogen. Disruption of these pathways by deletion of *phoPR* (Fig. 8), interruption of the *phoPR*-regulated gene *pks2* (Fig. 1), inhibition of PhoPR-by treatment with ETZ (Fig. 5), inhibition of *tgs1* expression by treatment with MSU-39446 (Fig. 8), or the absence of vitamin B12, results in the enhanced acid growth phenotype. Notably, treatment with PZA also leads to enhanced acid growth, suggesting part of PZA function may be related to inhibiting lipid synthesis. When lipid synthesis is blocked, growth on propionate at acidic pH is dependent on the MCC, which is one of primary mechanisms by which Mtb detoxifies propionate (38). Pyruvate is a permissive carbon source at acidic pH, therefore, it is expected that if propionate is metabolized to pyruvate via the MCC that it would promote growth (Figure 8). Given the many regulatory functions of PhoPR(39), it remains possible that PhoPR is also regulating lipid-synthesis independent mechanisms to arrest growth. Given the link between growth arrest and decreased CoA-pools, this could possibly be a regulatory activity inhibiting central metabolism. We have also not explored the potential role of cAMP signaling in the observed phenotypes, where PhoPR modulation of cAMP could be impacting propionate detoxification mechanisms and cell envelope permeability(40, 41).

**Figure 8:**
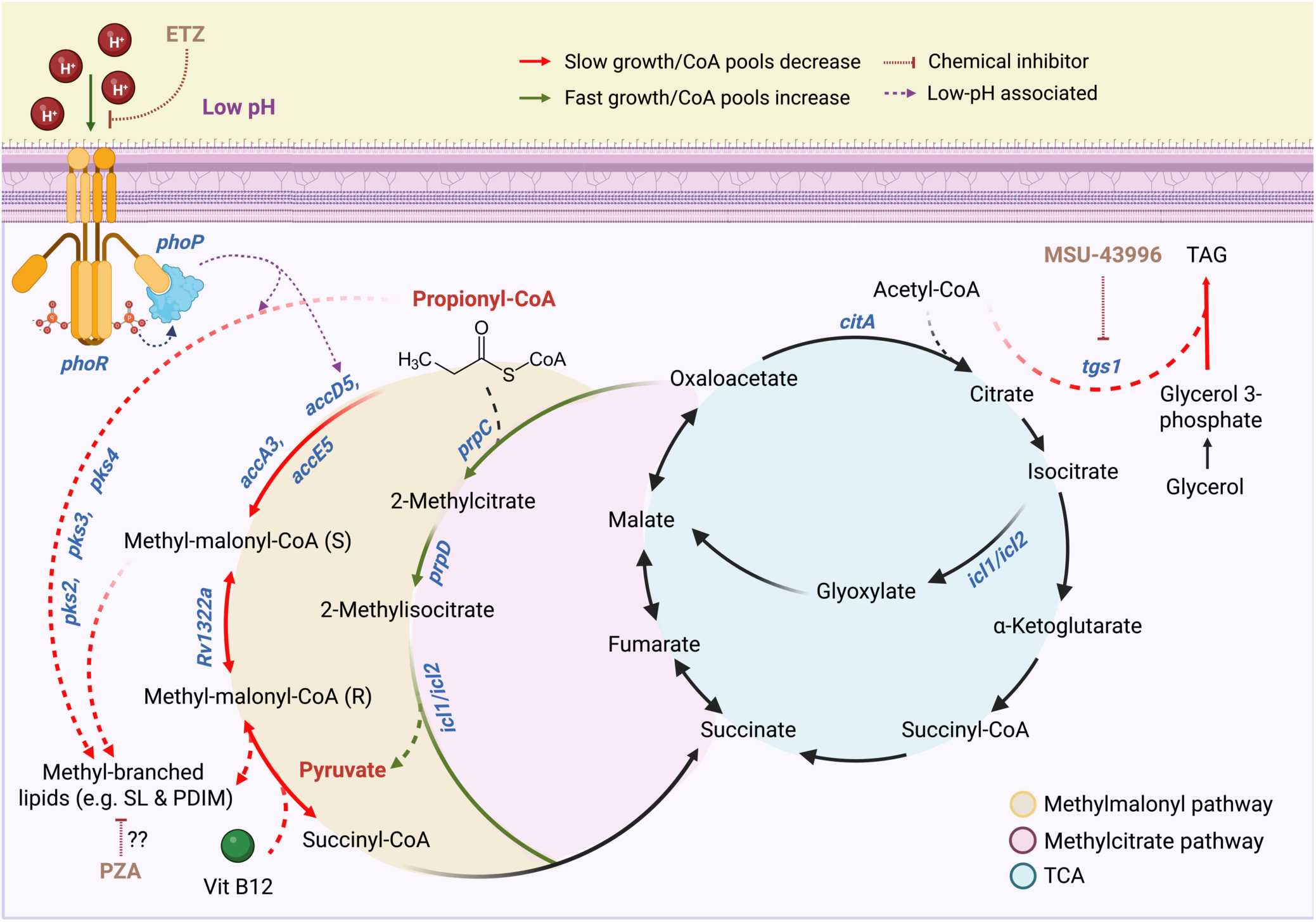
Model for Mtb growth arrest on propionate at acidic pH and mechanisms of suppression by mutations in *phoPR* or pyrazinamide treatment. Propionate-dependent acid growth arrest is a PhoPR-regulated response driven by diversion of carbon away from central metabolism and into the synthesis lipids. At acidic pH, propionyl-CoA is used as a substrate to synthesize methyl-branched lipids, either through the polyketide synthases Pks2, Pks3 and Pks4, or through the methyl-malonyl pathway. Additionally, the enzyme MutAB, a vitamin B12-dependent enzyme, catalyzes the reversible interconversion between malonyl-CoA and succinyl-CoA, contributing to the synthesis of branched lipids. In either cases, low pH triggers remodeling of central metabolism towards complex lipids synthesis to alter the mycomembrane structure as a pH-dependent adaptation, consuming the CoA pools and slowing, or arresting growth in the process. This pH-dependent adaptation in lost in the *phoPR* null mutant or under indirect PhoPR chemical inhibition using ethoxzolamide (ETZ). Instead, the propionate is metabolized into pyruvate by the methylcitrate cycle, a permissive carbon source, increasing CoA pools and permitting fast growth. Additionally, at acidic pH, the *phoPR* null mutant upregulates triacylglycerol synthesis using tgs1. Chemical inhibition of *tgs1* using MSU-39446 prevents diversion of CoA pools into TAG synthesis and away from central metabolism, increasing CoA pools, metabolism and growth in both the WT and *phoPR* null mutant. PZA treatment causes enhanced growth in the WT, which may be driven by inhibiting lipid synthesis via depletion of CoA pools.

The presented data show that the PhoPR plays a role in the detoxification of propionate. We observed that the DMSO treated WT strain and the single Δ*icl1* or Δ*icl2* mutants do not have a survival defect (Figure. 5 and S2) and the double Δ*icl1/2* mutant has its viability reduced by ∼1 log relatively to the starting CFUs. In contrast, when PhoPR signaling is inhibited by ETZ, viability is reduced in Δ*icl2* mutants and in the Δ*icl1/2* double mutant by ∼100 fold (Figure. 5 and S2). This demonstrates that inhibition of PhoPR sensitizes Mtb to propionate toxicity. Therefore, PhoPR is exerting dual functions under the tested conditions, to both arrest growth and detoxify propionate. Mtb induces PhoPR at acidic pH on various carbon sources (glycerol, pyruvate, propionate), and WT Mtb has a functional MCC, therefore, we do not believe that PhoPR is solely induced specifically to detoxify propionate. Rather, we observed that the PhoPR mutant experiences reductive stress at acidic pH, and we hypothesize that lipid synthesis serves the purpose of oxidizing cofactors (such as NADPH to NADP+) to maintain redox homeostasis(17, 18). Consistent with this hypothesis, we observe that downregulation of SL production in the *phoPR* mutant is associated with induction of other lipids such as TAG (17). Blocking both PhoPR and DosRST pathways removes four of the major lipid synthesis pathways (SL, DAT, PAT and TAG) and results in the most robust growth of propionate (Figure 5C, S2C). Pyruvate is known to promote growth at acidic pH, and we hypothesize that by entering the TCA cycle near the anaplerotic node provides Mtb the metabolic flexibility to maintain redox homeostasis by balancing anabolism and catabolism, possibly via the PEP glyoxylate cycle(42, 43). In this manner, if propionate is not diverted to lipid synthesis, it can be turned into a permissive carbon source that balances redox homeostasis at the anaplerotic node, even in the absence of *phoPR*.

Similar to the mutations in *phoPR*, PZA at low concentrations can also suppress Mtb growth arrest on propionate at acidic pH. Based on our model for propionate-dependent growth arrest, we reason that PZA treatment may be acting to inhibit lipid synthesis, thus enabling more propionate to be metabolized by the MCC into central metabolism. The mechanism of action of PZA remains controversial, but recent evidence supports a role for CoA metabolism and changes in cell envelope lipid synthesis playing a role in its activity (30, 44) . Resistance mutations to PZA arise in PanD, which is involved in the synthesis of CoA. The PZA metabolite POA can bind to PanD to inhibit CoA synthesis (32). Reduced CoA levels are predicted to result in lower levels of lipid synthesis, as it is an essential cofactor in the initial steps of fatty acid activation. Notably, it was recently reported that PDIM genes are strongly downregulated by treatment with PZA (45) and PDIM mutants are associated with resistance to PZA (44). Therefore, we speculate enhanced growth of the WT is due to reduced PDIM synthesis or other long-chain fatty acids that require CoA for their synthesis. The small range of concentrations where this enhanced growth in the WT is observed (∼3 µM) may reflect a hypomorphic phenotype, where CoA metabolism is sufficiently repressed by PZA to impact lipid metabolism, but not so much so to fully deplete CoA and kill the cell. It is surprising that the *ΔphoPR* mutant is susceptible to PZA and does not have further enhanced growth. Furthermore, the *ΔphoPR* mutant is more susceptible to PZA on both propionate and pyruvate (Fig. 7 and Fig. S8), and INH on propionate (Fig. 7G), but rather associated with enhanced growth at acidic pH. Because propionate is metabolized to pyruvate by the MCC it is consistent with our model that PZA sensitivity would be similar on either propionate and pyruvate. The *ΔphoPR* is also more sensitive to INH on propionate (Fig. 7G), suggesting more general mechanisms be driving antibiotic susceptibility in the mutant. We speculate that an alternative mechanism is driving PZA or INH sensitivity in the *phoPR* mutant, possibly differential expression of a *phoPR* pathway independent on lipid anabolism. For example, a *ppe51* loss of function mutant is reported to have enhanced susceptibility to PZA(46). *ppe51* is downregulated in the *phoPR* mutant, suggesting it, or some other unknown PhoPR*-*regulated genes may play a role in susceptibility that overcomes the expected further enhanced growth. Alternatively, the *phoPR* mutant may have has increased permeability due to changes in the cell envelope or cAMP signaling (13, 40, 47). It is possible that this increased permeability is increasing the intracellular concentration of PZA or INH, accumulating to sufficient levels to inhibit growth.

### Concluding remarks

The observation that PhoPR restricts growth on propionate at acidic pH is analogous to the findings of Baek *et al*, who showed that the DosRST and hypoxia regulated gene *tgs1* inhibits growth during hypoxia by diverting carbon from the TCA cycle(26). In both cases, growth restriction is an environmental adaption and not an inherent metabolic limitation. TB disease is caused by slow growing mycobacteria and persistence and growth arrest are associated with drug tolerance. By understanding the complex mechanisms by which Mtb integrates environmental cues, metabolism and growth arrest, we may define new drug targets that can disrupt pathogenesis in a way the increases bacterial growth. Enhanced growth is associated with reduced fitness in macrophages (36) and enhanced drug susceptibility *in vitro* (18). Therefore, these non-traditional drug targets may have unique impacts during treatment, including treatment shortening. Indeed, PZA was essential to shortening TB treatment times to six months and it is tempting to speculate that some of this activity may be related to PZA enhancing Mtb growth in mildly acidic and cholesterol rich environments, such as the macrophage or the granuloma. It is also important to consider that *phoPR* and *prpR* have variants in clinical isolates (33–35), supporting that the interplay of PhoPR and propionate observed in these studies, maybe associated with differential pathogenesis and drug susceptibility and in human infections and drug treatment.

## Methods

### Bacterial strains and growth

Mtb experiments were conducted with WT ERDMAN or WT CDC1551 as indicated. The *ΔphoPR* deletion mutant, and complemented strains are in the CDC1551 background (10) and the *icl1/2* mutants and complemented strains are in the Erdman background(24). Cultures were maintained in 7H9 media supplemented with 10% OADC and .05% Tween-80. All single carbon source experiments were performed in MMAT minimal media buffered to acidic pH 5.7 or pH 7.0(21), using (1 g l−1 KH2PO4, 2.5 g l−1 Na2PO4, 0.5 g l−1 (NH4)2SO4, 0.15 g l−1 asparagine, 10 mg l−1 MgSO4, 50 mg l−1 ferric ammonium citrate, 0.1 mg l−1 ZnSO4, 0.5 mg l−1 CaCl2, and 0.05% Tyloxapol) . For 12-day growth experiments, time points were taken every three days. For each of our growth curve experiments, 2mM propionate was used unless stated otherwise. Chemical inhibitors and supplements were used at the following concentrations: Ethoxzolamide at 40 µM, MSU-39446 at 40 µM, Itaconic acid at 2 mM and vitamin B12 at 10ug/ml. Mtb was seeded in T-25 standing tissue culture flasks in 8ml of minimal media at an initial cell density of 0.05 OD and incubated at 37 °C and 5% CO_2_. 500 μL of each culture was taken each day for optical density measurements. Bacterial viability was assessed enumerating colony-forming units (CFUs) on 7H10+OADC plates.

### Genetic selection

Transposon mutagenesis was performed in Mtb Erdman using the ϕMycoMarT7 Tn system as previously described(48), generating a library with ∼100,000 Tn mutants. The Tn mutant library was plated on MMAT pH 5.7 agar plates supplemented with 2mM propionate as the sole carbon source. Plates were incubated at 37°C, with mutants appearing around week 8 and isolated for growth. Single-colony isolates were confirmed as EAG (enhanced acidic growth) mutants under acidic conditions in liquid media MMAT pH 5.7 supplemented with 2mM propionate. The transposon insertion sequences were confirmed using inverse PCR (49) or whole genome sequencing.

### Whole Genome Sequencing

Genomic DNA of selected mutants as well as the WT Erdman control was isolated, DNA libraries were constructed, and sequenced using the Illumina MiSeqm in-paired end, 250 bp read format (PE250). After the sequencing run, reads were demultiplexed and converted to FASTQ format using the illumina bcl2 fastq (v1.8.4) script. The reads in the raw data files were then subjected to trimming of low-quality bases and removal of adapter sequences using Trimmomatic v0.36, with a 4 bp sliding window, and a read quality cutoff of below 15 or read length cutoff less than 36 bp. The trimmed reads were then aligned to the Erdman reference genome using the Burrow-Wheeler alignments. Genome analysis tool kit base quality score recalibration, indel alignment, and duplicate removal were applied, and SNP and INDEL discovery were performed.

### pH and propionate dose response combination growth assays

Mtb cultures were incubated in a range of pH-buffered MMAT media ranging from pH 5.0-7.0 at a starting OD of 0.2 in 96 well plates. Cultures were treated with a 2.5-fold serial dilution (0mM-20mM) of propionate and incubated over a course of 21 days, with growth assessed by optical density.

### Coenzyme A assay

Coenzyme A was measured using a commercially available assay (Sigma-Aldrich). Cultures were maintained in 7H9, and then seeded to an OD of 0.2 and resuspended in minimal media supplemented with 2mM propionate in 8 ml of media. The assay measures both free CoA and acyl-CoA. Total CoA was measured at Day 0, 3, and 6 and concentration determined using a standard curve.

### Pyrazinamide Sensitivity Assays

Dose response assays were conducted in a 96 well plate cultures seeded at OD 0.2 and in minimal media supplemented with 2mM propionate across a dose response from 100 µM-0 µM of PZA. Plates were incubated for 6 days in a sealed Ziploc bag at 37C and OD read on a plate reader. Standing flask assays were conducted using conditions as described earlier in the methods. WTCDC, *ΔphoPR* mutant, and complement were grown in minimal media at acidic pH supplemented with propionate and treated with DMSO, 3.84 uM Pyrazinamide, and 20uM Isoniazid using conditions described above. Growth was measured Day 0, 3, 6, 9, and 12. Viability was measured Day 0 and Day 12 by enumerating CFUs.

## Acknowledgements

We thank Prof. John McKinney for sharing the *icl1/icl2* mutant strains. Research on this project was supported by grants from the NIAID (R01 AI116605 and R01 AI150855) and AgBioResearch.

## Supplemental Table and Figures

### Supplemental Table

**Table S1:**
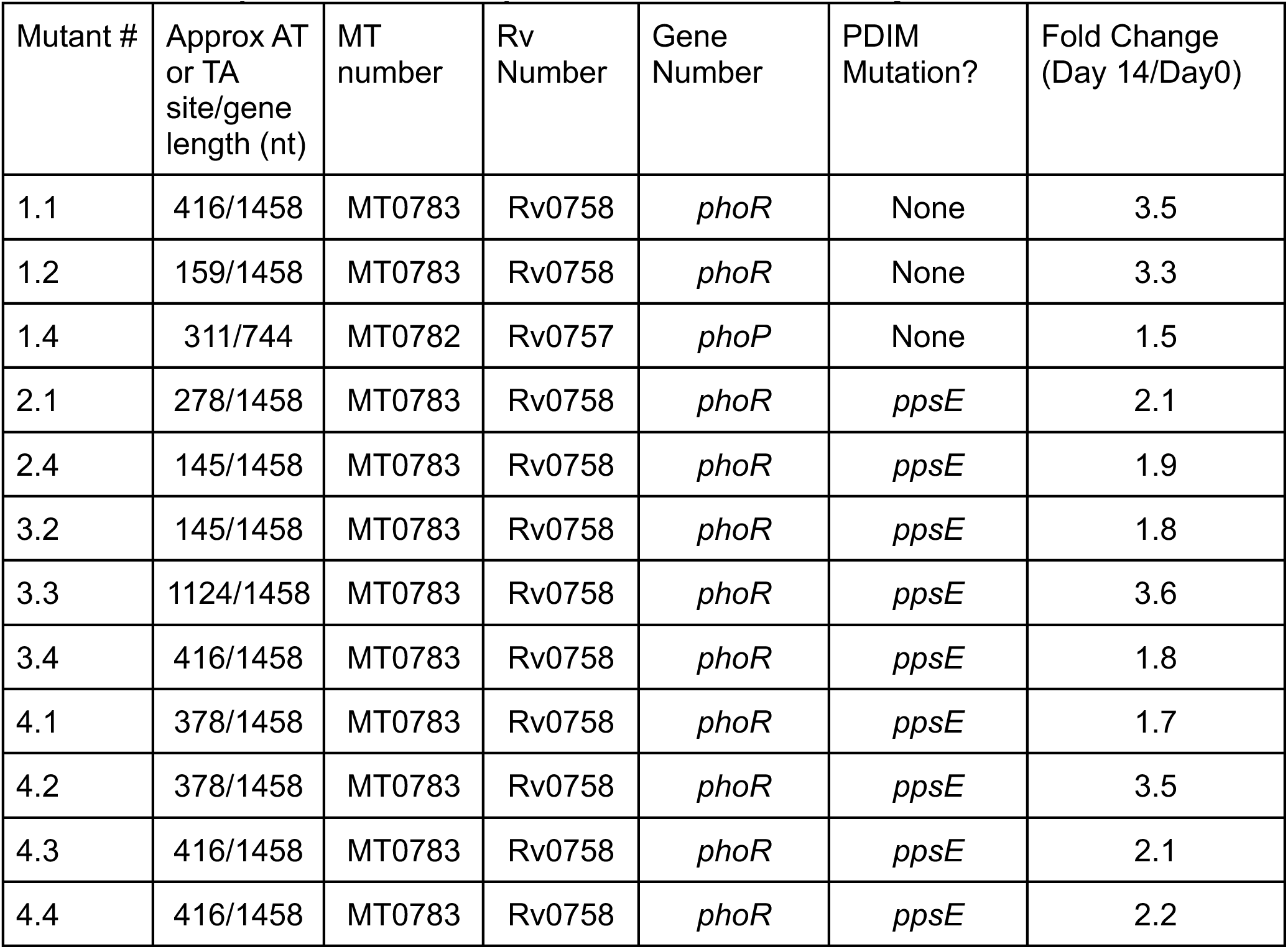
Transposon Mutant Propionate Selection Summary.

### Supplemental Figure Legends

**Figure S1.**
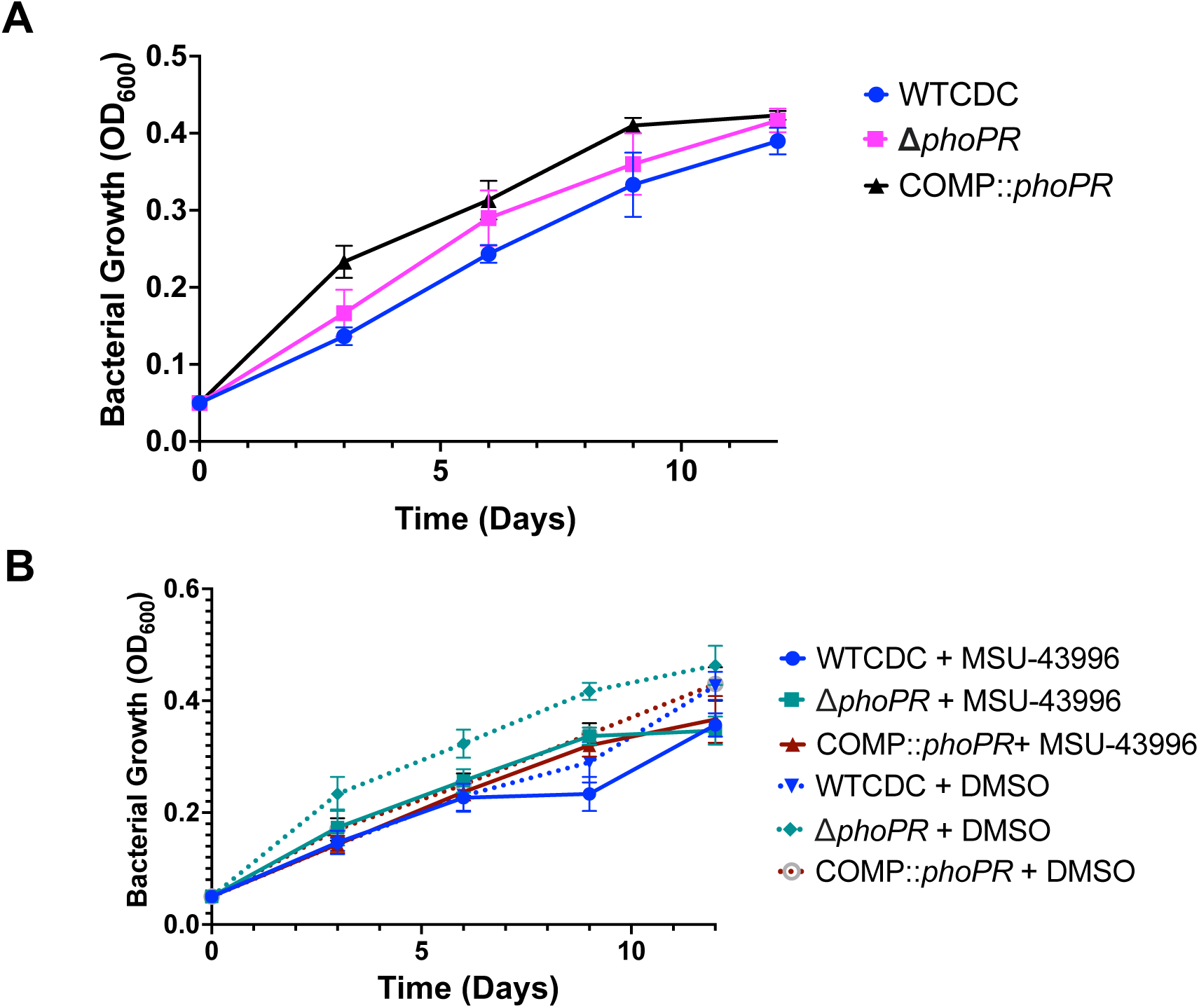
Bacterial Growth and viability of WT, τι*phoPR* mutant, and complemented strains grown on propionate and treated with MSU-43996 at neutral pH. A) bacterial growth of the three strains on propionate at pH 7.0, showing no significant differences in growth rate. B) bacterial growth of the three strains on propionate at pH 7.0, treated with MSU-43996 or the vehicle, showing no major differences in growth rates based on strain or treatment.

**Figure S2.**
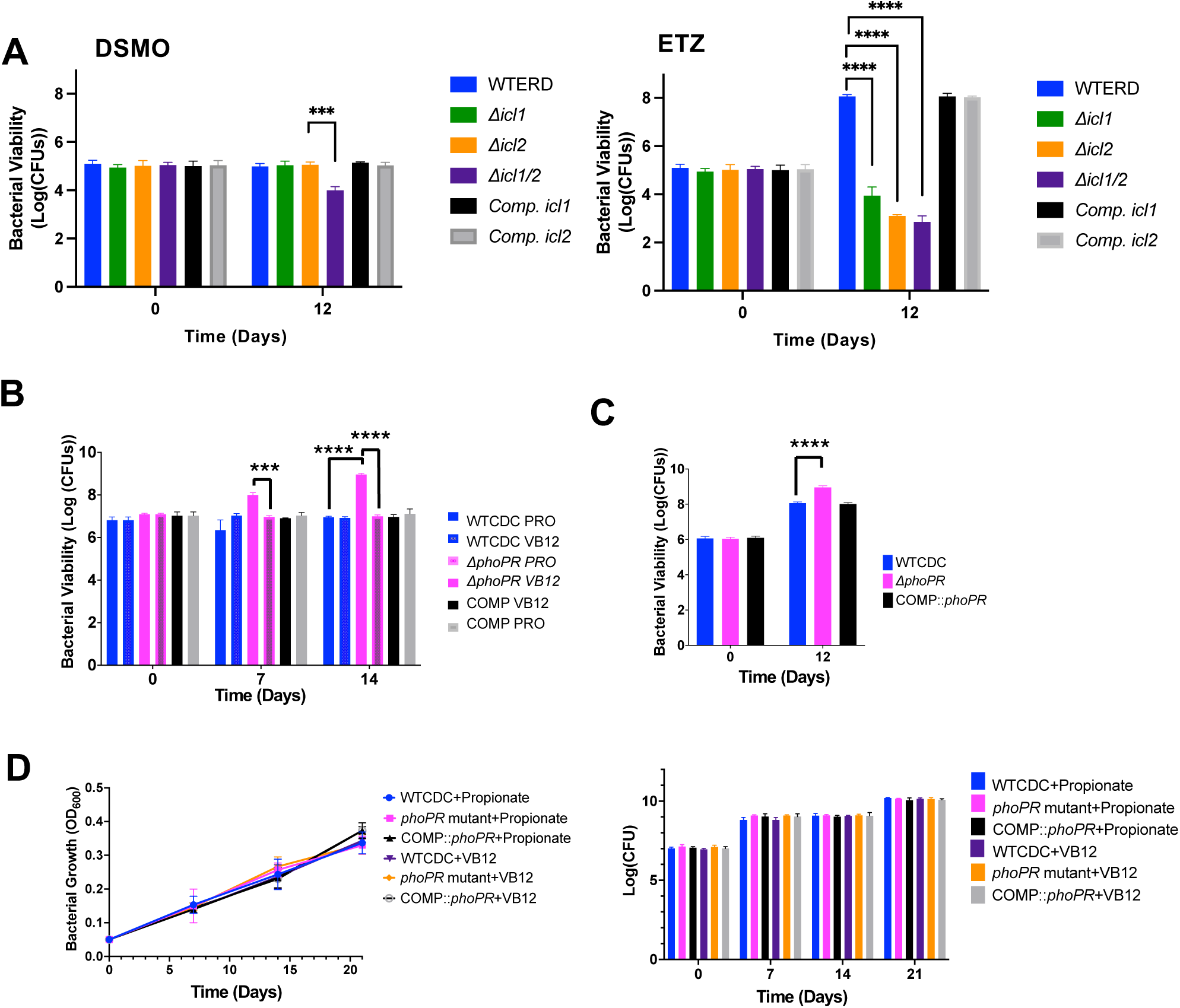
PhoPR restricts growth at acidic pH by diverting carbon from the methyl citrate cycle. **A)** CFUs from the corresponding experiment in Figure 5A, showing cell death in the ETZ treated icl2 and icl1/2 knockouts. **B)** CFUs of Vitamin B12 treated Mtb strains. **C)** CFUs for figure 5C showing enhanced growth of MSU-43996 treated Mtb. Note, that timepoints are different, but CFUs are from the same experiment. **D**) Vitamin B12 does not impact the growth of strains at neutral pH. A one-way ANOVA was used for the viability experiments, **\*** <0.05, **\*\*** <0.01, **\*\*\*** <0.001, **\*\*\*\*** <0.0001. Experiments were replicated at least twice with similar results.

**Figure S3.**
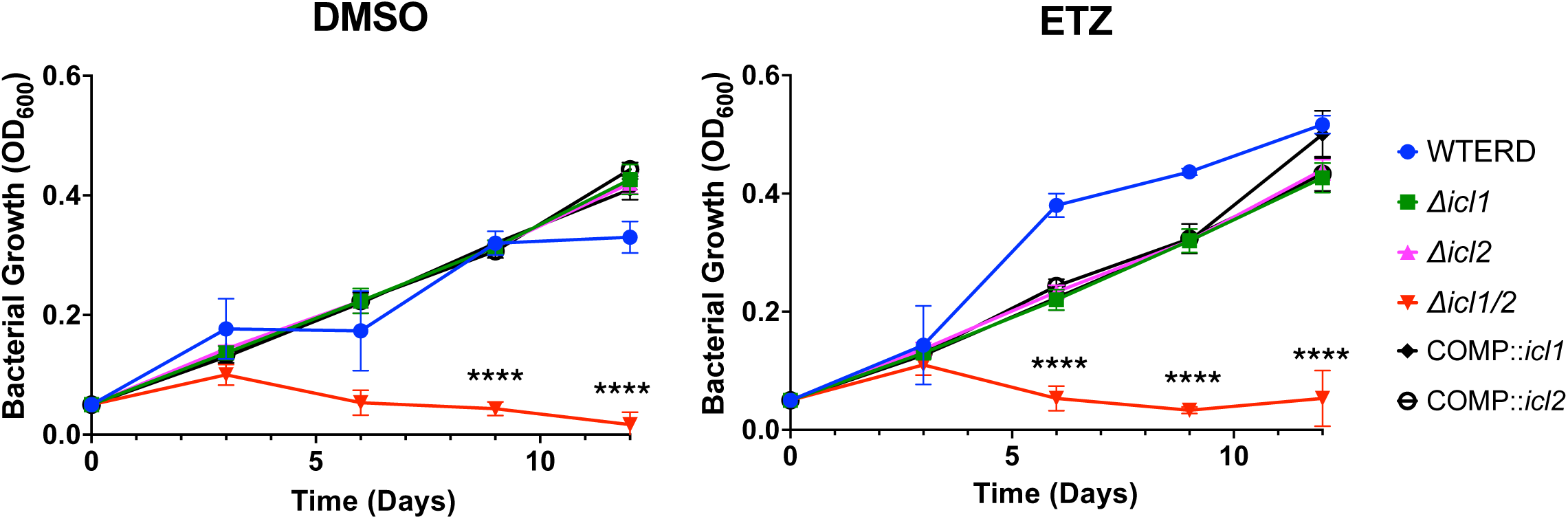
Function of Icl and PhoPR at neutral pH on propionate. Growth of Mtb strains at pH 7.0 on propionate. ETZ did not impact growth in the *Icl1* or *Icl2* single mutants. The double icl1/2 mutant dies due to propionate toxicity. An unpaired t-test was used between individual groups for the growth curves, **\*** <0.05, **\*\*** <0.01, **\*\*\*** <0.001, **\*\*\*\*** <0.0001. Experiments were replicated at least twice with similar results.

**Figure S4.**
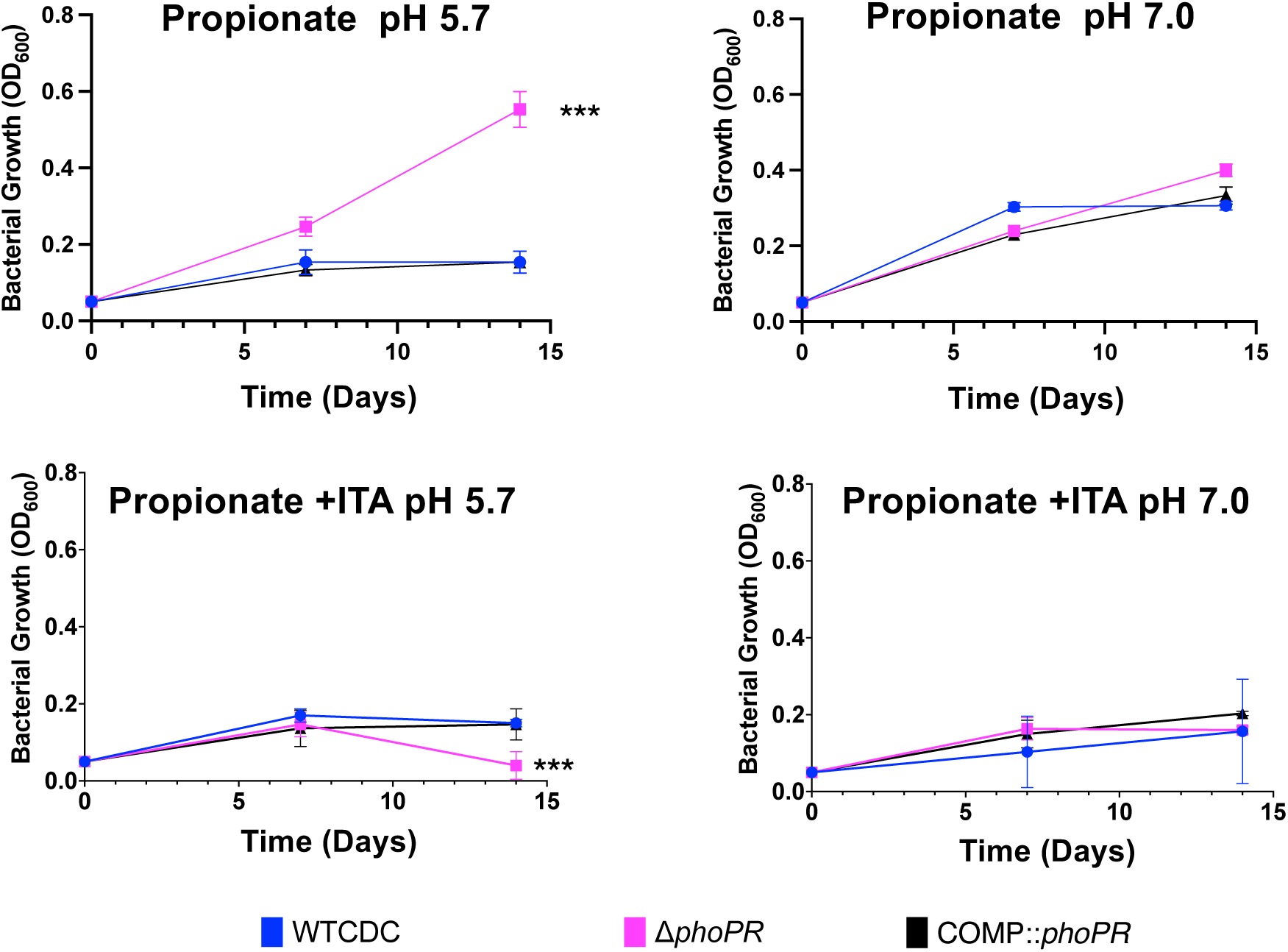
Itaconic acid inhibits enhanced growth of the Δ*phoPR* mutant. Itaconic acid inhibits the ability of the Δ*phoPR* to grow on propionate at pH 5.7. An unpaired t-test was used between individual groups for the growth curves, **\*** <0.05, **\*\*** <0.01, **\*\*\*** <0.001, **\*\*\*\*** <0.0001. Experiments were replicated at least twice with similar results.

**Figure S5.**
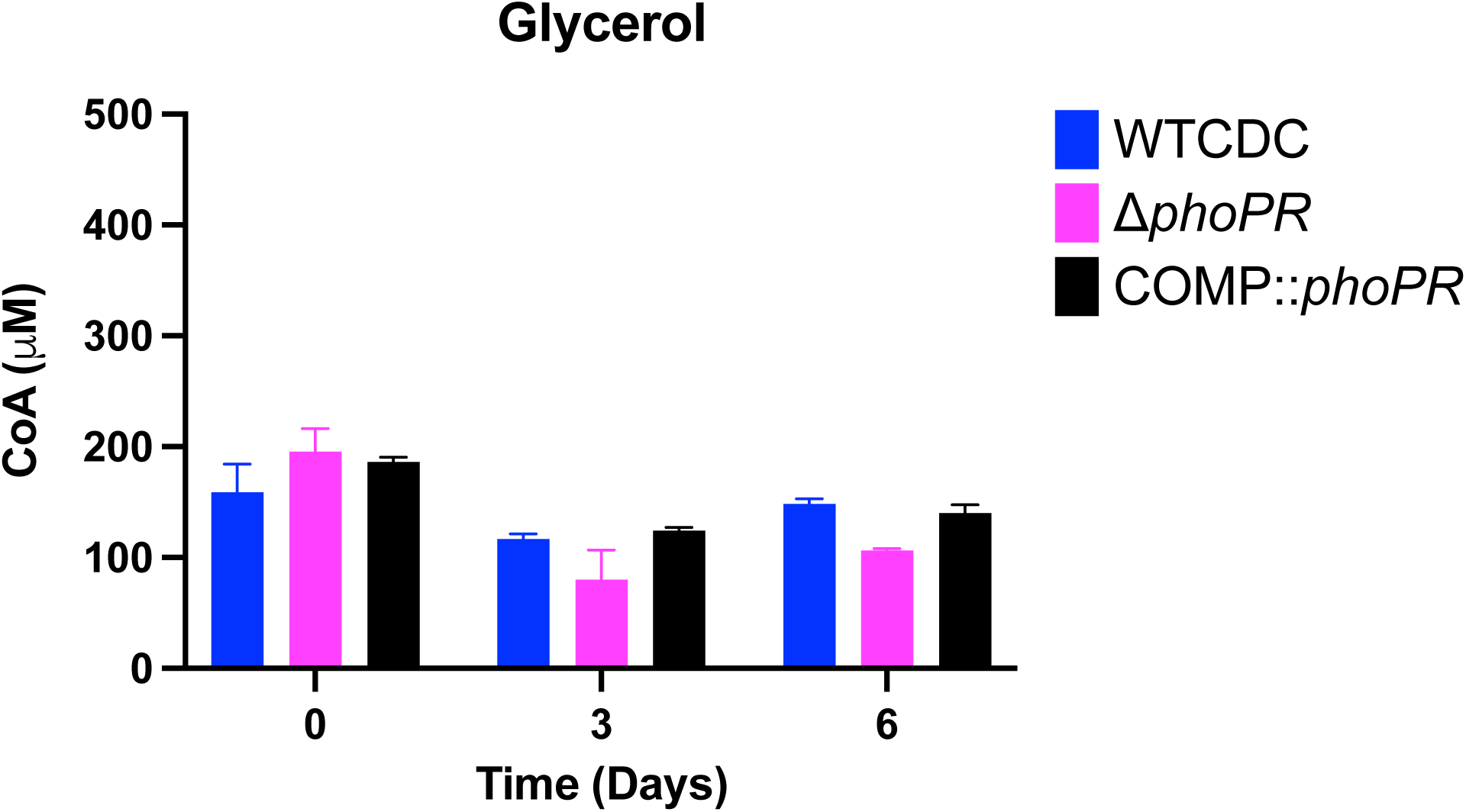
Free CoA pools on glycerol as a sole carbon source. During acid growth arrest on glycerol, all three strains maintain relatively low free CoA pools. Two-way ANOVA was used for this analysis, **\*** <0.05, **\*\*** <0.01, **\*\*\*** <0.001, **\*\*\*\*** <0.0001. Experiments were replicated at least twice with similar results.

**Figure S6.**
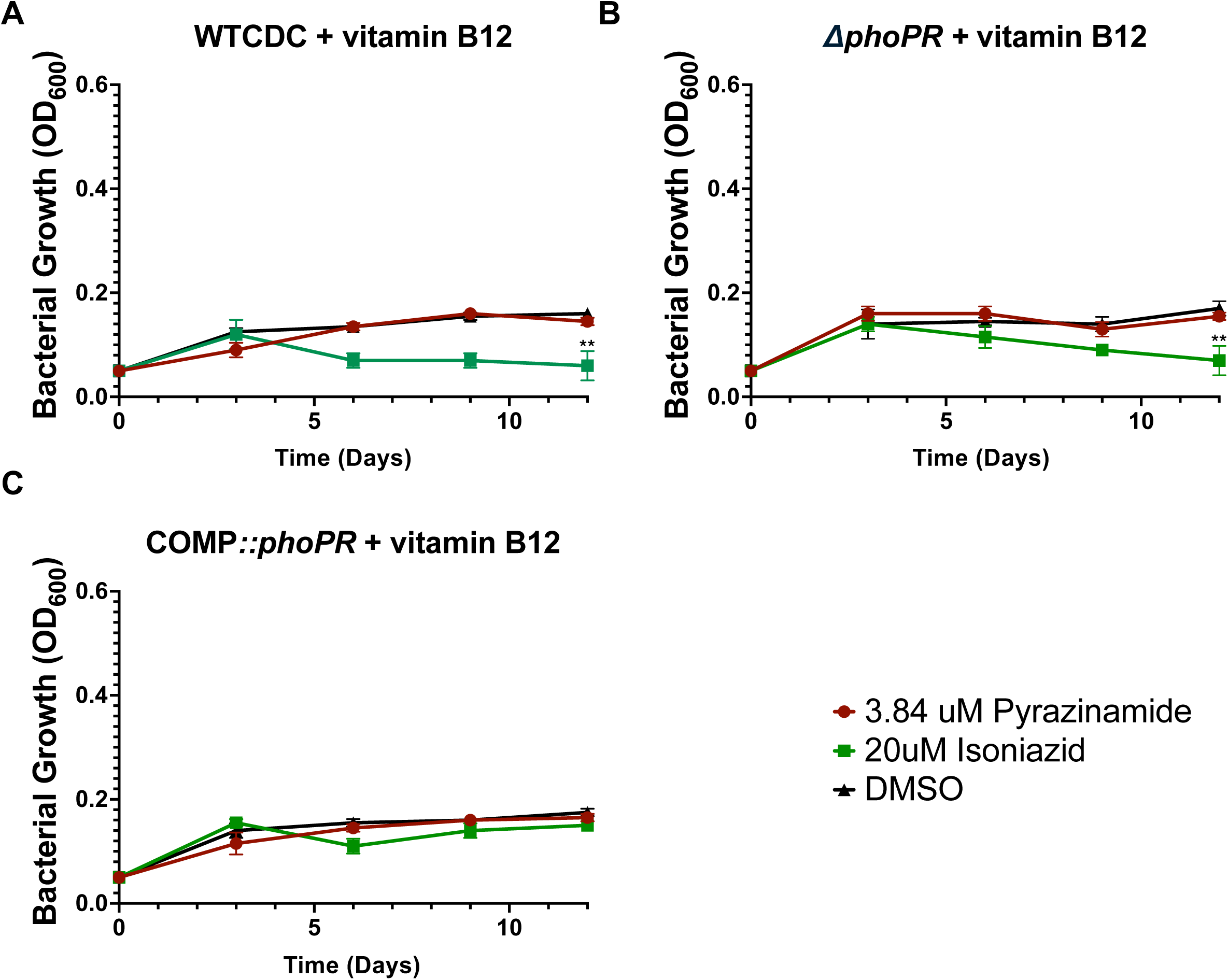
Vitamin B12 inhibits enhanced growth of WT Mtb treated with PZA. The inhibition of enhanced growth is consistent with PZA inhibiting lipid synthesis to promote growth, as making another lipid synthesis pathway available abolishes enhanced growth. Multiple comparison unpaired t-test was used for the analysis between treatments and the control, and between strains receiving the same treatment, **\*** <0.05, **\*\*** <0.01, **\*\*\*** <0.001, **\*\*\*\*** <0.0001. Experiments were replicated at least twice with similar results.

**Figure S7.**
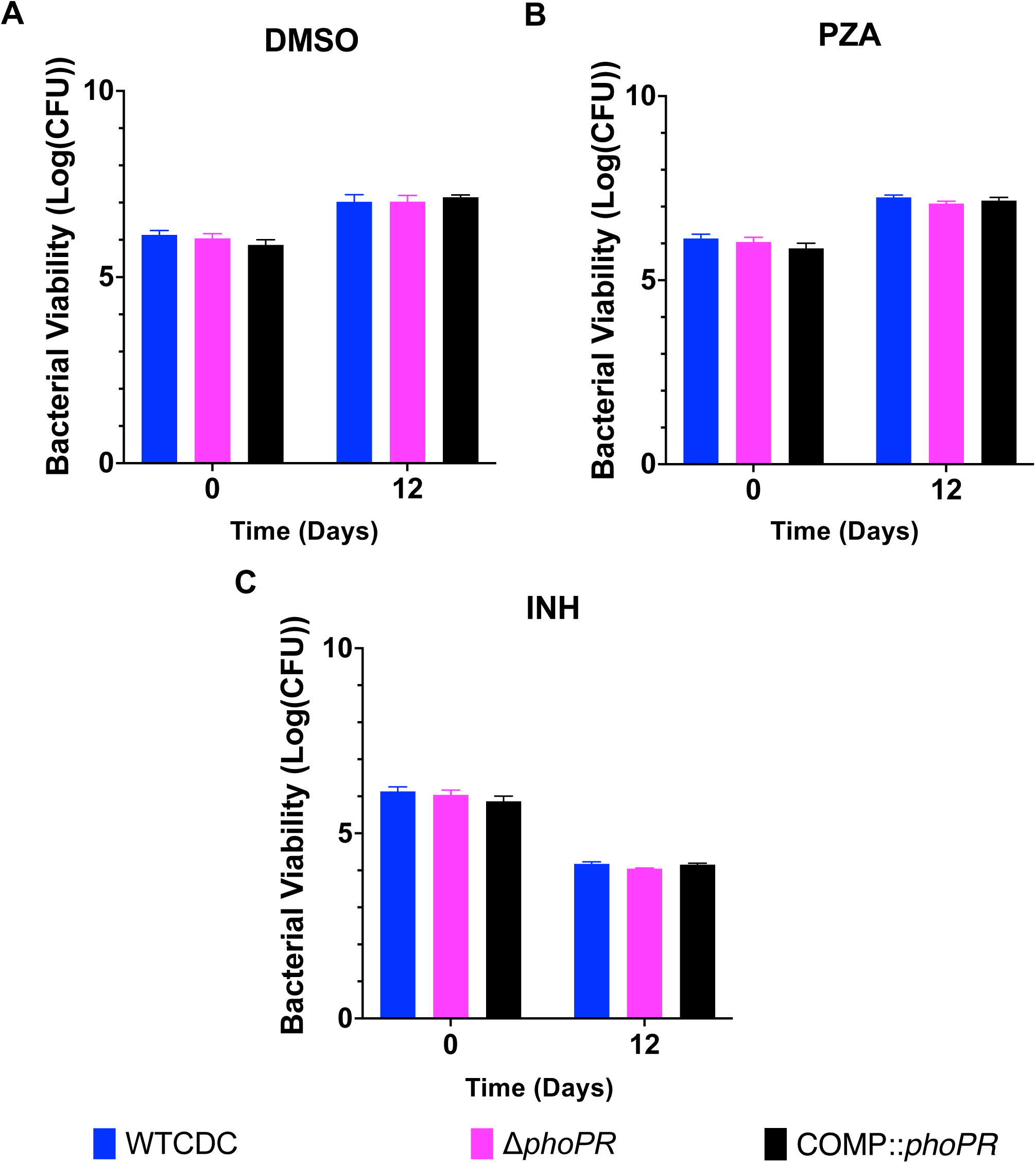
Vitamin B12 inhibits enhanced growth of WT Mtb treated with PZA. CFU data for OD data presented in Figure S6.

**Figure S8.**
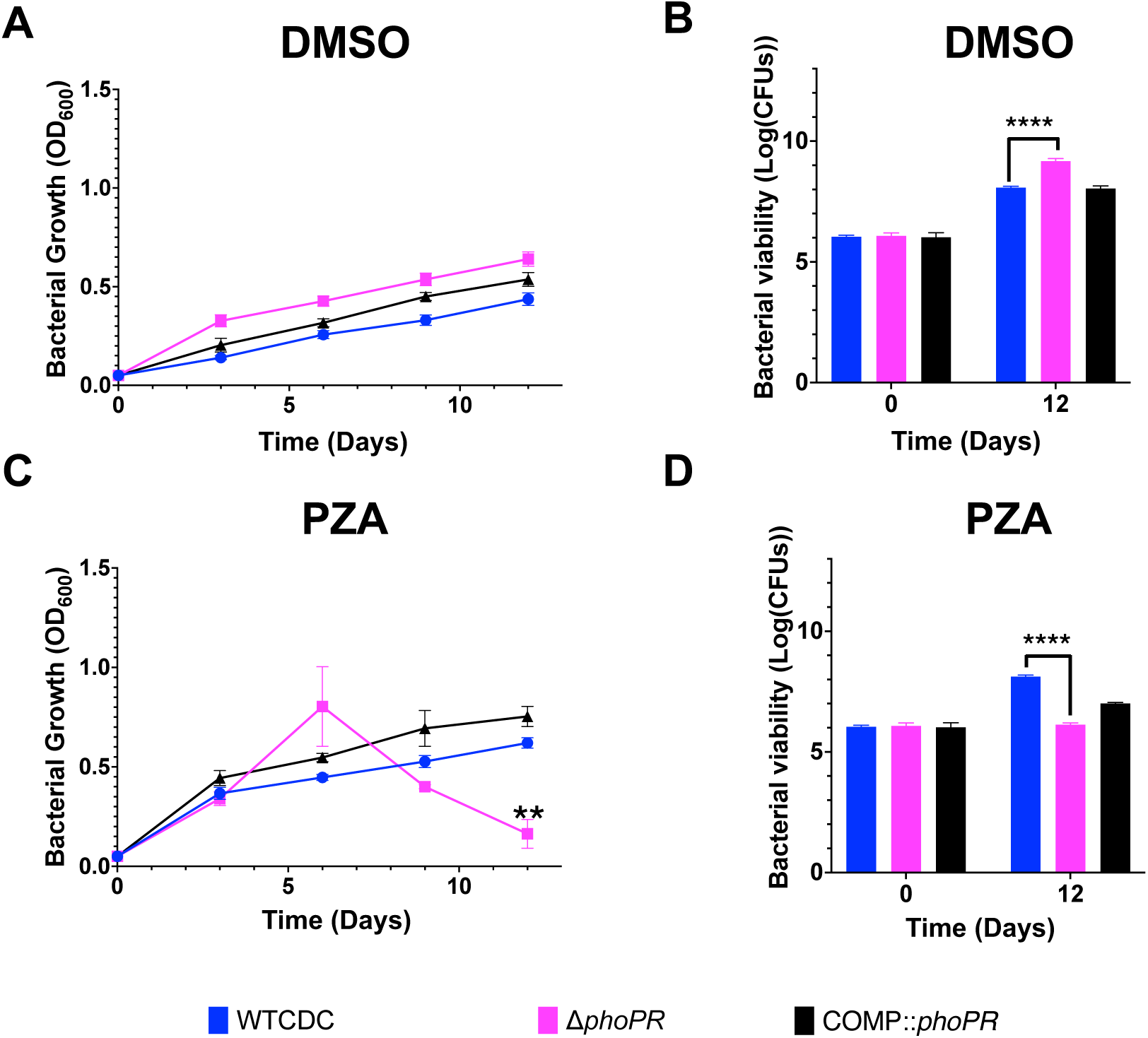
Sensitivity of Mtb to PZA on Pyruvate as a sole carbon source at pH 5.7. PZA treatment does not result in enhanced growth with pyruvate as a sole carbon source.

**Figure S9.**
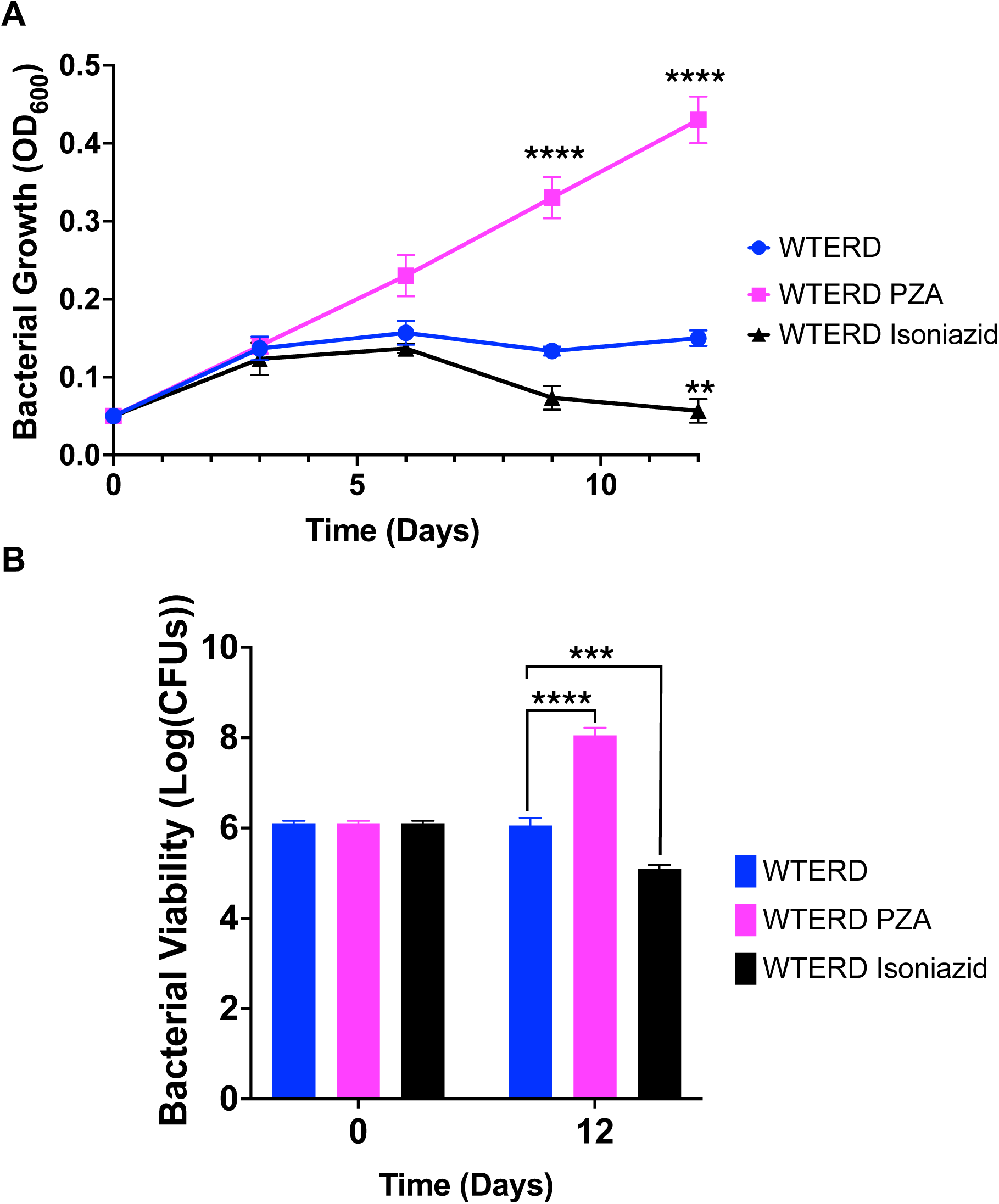
PZA suppresses acid growth arrest on propionate in Mtb Erdman. To determine if the enhanced growth by PZA is conserved across strains, the PZA treatment experiment from Figure 7 which as conducted in CDC1551, was repeated in Mtb Erdman. Multiple comparison Unpaired t-tests were used for the growth assays, and one-way ANOVA was used for the viability assays, **\*** <0.05, **\*\*** <0.01, **\*\*\*** <0.001, **\*\*\*\*** <0.0001. Experiments were replicated at least twice with similar results.

**Figure S10.**
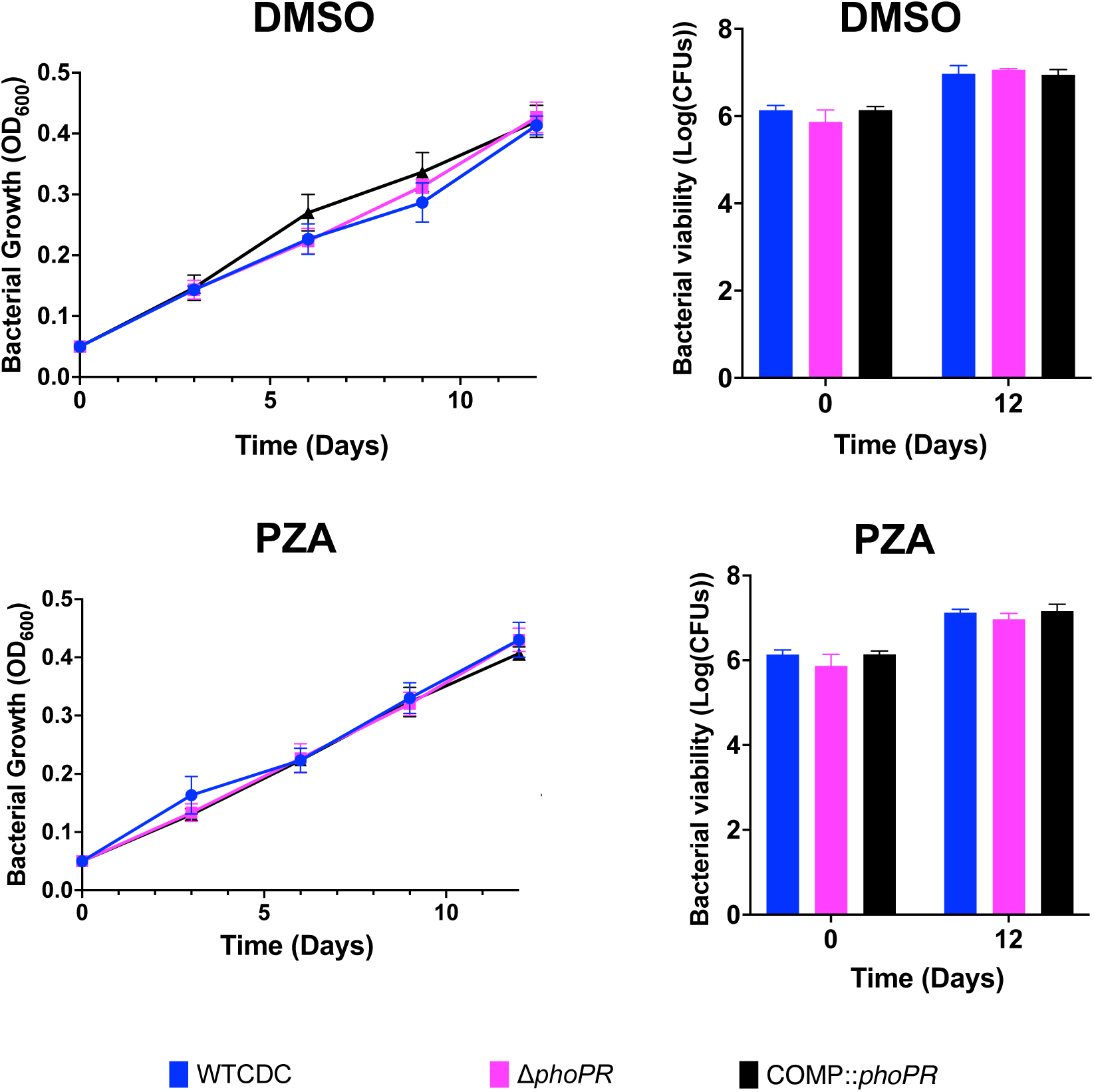
Sensitivity of Mtb to PZA treatment at pH 7.0. Control experiment for data in Figure 7, showing PZA is not active on Mtb grown in propionate at pH 7.0.

## Notes

### Summary of Updates

The manuscript is revised based comments from peer review. There are no substantive changes to the figures or conclusions. The minor changes include: 1) Updates to Figure 5A, Figure 6 and Table S1, Supplementary Figure 2, Supplementary Figure 5. 2) We added some clarifying statements to the discussion. 3) We added a middle initial for the first author.

